# Comparative evaluation of cell-based assay technologies for scoring drug-induced condensation of SARS-CoV-2 nucleocapsid protein

**DOI:** 10.1101/2024.09.26.615262

**Authors:** Rui Tong Quek, Cyna R. Shirazinejad, Christina L. Young, Kierra S. Hardy, Samuel Lim, Phillip J. Elms, David T. McSwiggen, Timothy J. Mitchison, Pamela A. Silver

**Author notes:** Correspondence (T.J.M.), (P.A.S.).

## Abstract

Protein-nucleic acid phase separation has been implicated in many diseases such as viral infections, neurodegeneration, and cancer. There is great interest in identifying condensate modulators (CMODs), which are small molecules that alter the dynamics and functions of phase-separated condensates, as a potential therapeutic modality. Most CMODs were identified in cellular high-content screens (HCS) where micron-scale condensates were characterized by fluorescence microscopy. These approaches lack information on protein dynamics, are limited by microscope resolution, and are insensitive to subtle condensation phenotypes missed by overfit analysis pipelines. Here, we evaluate two alternative cell-based assays: high-throughput single molecule tracking (htSMT) and proximity-based condensate biosensors using NanoBIT (split luciferase) and NanoBRET (bioluminescence resonance energy transfer) technologies. We applied these methods to evaluate condensation of the SARS-CoV-2 nucleocapsid (N) protein under GSK3 inhibitor treatment, which we had previously identified in our HCS campaign to induce condensation with well-defined structure-activity relationships (SAR). Using htSMT, we observed robust changes in N protein diffusion as early as 3 hours post GSK3 inhibition. Proximity-based N biosensors also reliably reported on condensation, enabling the rapid assaying of large compound libraries with a readout independent of imaging. Both htSMT and proximity-based biosensors performed well in a screening format and provided information on CMOD activity that was complementary to HCS. We expect that this expanded toolkit for interrogating phase-separated proteins will accelerate the identification of CMODs for important therapeutic targets.

## Introduction

Biomolecular condensates are dynamic assemblies containing dense networks of protein and nucleic acid molecules and are important for many cellular functions^1–3^. They form by liquid-liquid phase separation often driven by weak multivalent protein-protein and protein-nucleic acid interactions, which allows for rapid assembly and dispersion upon changes in environmental stimuli. Functionally, these condensates spatially organize cellular compartments to accelerate biochemical reactions and to sequester molecules for regulation of cell signaling^3^. Several examples of biomolecular condensates include the nucleolus for ribosomal RNA synthesis and ribosome assembly^4^, P granules for germ cell specification^5^, and stress granules for protective sequestration of RNAs upon various stresses^6^.

Given the importance of biomolecular condensates to cellular functions, dysregulation of condensation has been implicated in numerous diseases^7–9^. In the context of neurodegenerative diseases like amyotrophic lateral sclerosis (ALS) and frontotemporal dementia (FTD), genetic mutations in RNA-binding proteins like FUS and hnRNPA1 promote phase transitions and the formation of amyloid-like fibrils^10, 11^. Genetic mutations also result in abnormal condensation of the alternative splicing factor RBM20 in congenital dilated cardiomyopathy^12^, and in some cancers, these mutations can lead to the formation of aberrant transcriptional condensates that upregulate oncogenes^13, 14^. Furthermore, the formation of *de novo* viral condensates in infected cells is critical to various steps of viral lifecycles in viruses such as Influenza A, SARS-CoV-2, and *Mononegavirales*^15–24^. As such, evaluation of small molecule condensate modulators (CMODs) that modify condensate components or dynamics has become increasingly of interest. In particular, CMODs that target disease-driving condensates have emerged as an important therapeutic modality^25–27^. Several examples include CMODs targeting pathogenic RSV viral condensates^28^, MED1 and BRD4 nuclear condensates^29^, stress granules^30^, and fusion oncoprotein condensates^31^ (among many others), demonstrating the importance of investigating these small molecules and their role in addressing disease. Therefore, there is a significant need for drug discovery tools to robustly identify therapeutically relevant CMODs.

To date, most CMODs were identified by high-content screens (HCS) in which condensing target proteins were imaged in cells, with or without fixation. In such screens, CMOD activity was detected by changes in condensate properties such as number, size, and morphology, following image analysis^30, 32–35^. For example, we screened for CMODs targeting the SARS-CoV-2 nucleocapsid (N) protein by stably expressed N-EGFP in human cells^32^ and identified compounds that modulated N condensation from a library of bioactive molecules. Our assay involved seeding cells, treating cells with a small molecule library for 24h, fixing cells, and finally staining with dyes before imaging with an automated fluorescence microscope (Figure 1A). Image analysis was performed to identify and count the number of puncta per cell under each treatment condition (Figure 1B). We found that ATP-competitive GSK3 inhibitors induced condensate formation and hardening of multiple coronavirus N proteins. Consistent with on-target action, the EC_50_ values of multiple structurally diverse inhibitors for N condensation paralleled the EC_50_ values for Wnt pathway induction, a pathway known to be modulated by GSK3. We showed that both the ATP-competitive GSK3 inhibitors LY2090314 and CP21R7 induced N condensation, with LY2090314 being more potent than CP21R7, whereas the allosteric GSK3 inhibitor tideglusib was inactive in inducing N condensation (Figure 1B).

**Figure 1:**
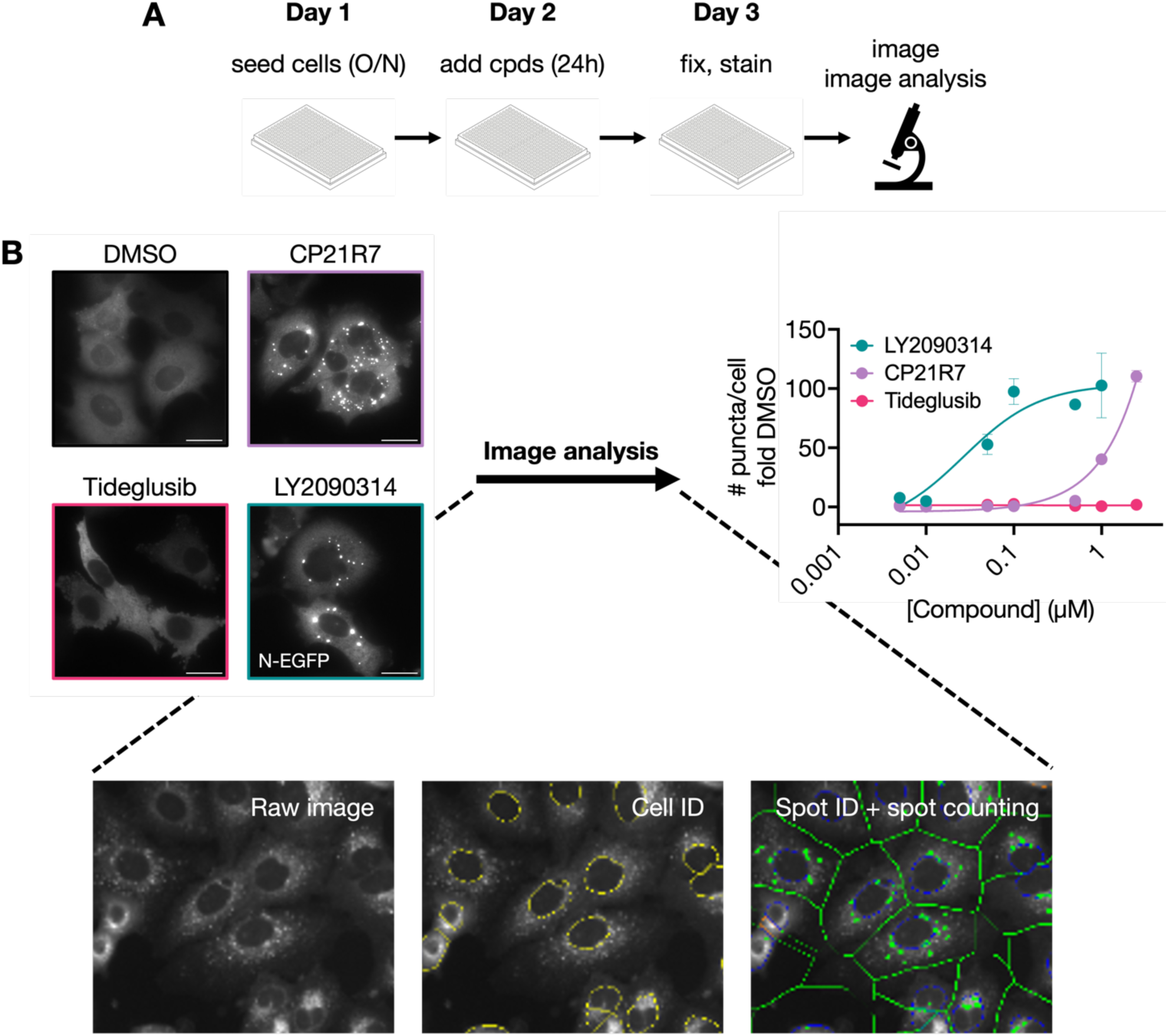
High-content screening (HCS) assay set-up and drawbacks for identifying small molecule protein condensate modulators (CMODs). *A.* Workflow for identifying CMODs that induce SARS-CoV-2 N protein condensation by high content imaging. O/N: overnight; cpds: compounds. *B.* HCS image analysis workflow for characterizing CMOD effects on N condensation states. Left: fluorescence images of A549 cells stably expressing N-EGFP, upon 24h treatment with 10µM GSK3 inhibitors. Scale bar = 20µm. Bottom: an example image analysis pipeline for spot counting and quantification for each image acquired during the screen. Right: Bar graph illustrating the number of puncta (condensates) per cell as a fold change over DMSO control. Adapted from Quek et. al., 2023^32^.

While we observed robust condensate formation for select compounds, this HCS assay had several drawbacks. HCS assays struggle with throughput and resolution – imaging a plate with sufficient coverage takes much longer than reading out a plate with a plate reader-compatible signal. In addition, HCS suffers from optical resolution limitations, only allowing the differentiation of condensates above the microscope diffraction limit. This limits the characterization of smaller condensates such as transcriptional condensates. While increasing imaging resolution is possible with super resolution approaches, these instruments lack the throughput needed for assaying large chemical libraries. Furthermore, image analysis pipelines for condensate characterization are overfit for pre-defined condensate phenotypes and are sensitive to artifacts from non-specific fluorescence background. Finally, many of these HCS campaigns are performed with a single snapshot of cells after compound treatment, failing to resolve condensate dynamics that may capture information averaged out at a single time point.

To increase speed, reliability, and information content of CMOD characterization, we investigated two alternative technologies that are compatible with high-throughput screening – fast high-throughput single molecule tracking (htSMT) and proximity-based luciferase condensate biosensor assays. SMT is a super resolution technique used to follow protein motion to investigate various biological phenomena^36^. However, traditional SMT techniques have been incompatible with large scale high-throughput screening due to unreliable data quality and coverage. htSMT technology combines a high-throughput imaging format with expanded imaging coverage to enable capturing protein motion and dynamics in a scalable format^37^. On the other hand, proximity-based biosensors based on split luciferase (NanoBIT) and bioluminescence resonance energy transfer (NanoBRET) technologies have become important drug discovery tools in recent years^38–42^. They have broad applicability in identifying conditions that promote or inhibit protein-protein interactions and can be used in living cells or in recombinant systems. These proximity-based biosensors report on nanometer-scale interactions; however, like htSMT, they have not yet become standard tools for discovering and characterizing CMODs. In both cases, these approaches allow the investigator to measure changes in the protein-protein interactions that give rise to condensate formation rather than the condensate end state, offering potential advantages in assay sensitivity as well as in driving mechanistic insight. We thus sought to employ both htSMT and proximity-based biosensors to studying CMOD modulation of SARS-CoV-2 N condensation as alternative CMOD identification and characterization strategies that do not rely on identification and quantification of N puncta.

We found that both htSMT and proximity-based biosensors reported on N condensation states in response to ATP-competitive GSK3 inhibitors robustly, with complementary advantages. Specifically, N condensation resulted in rapid and robust reduction in N protein diffusion when scored with htSMT, as well as increased proximity-induced signal in the NanoBIT and NanoBRET assays, all with dose dependent effects. These assays can be performed in a high-throughput context compatible with small molecule screening. Altogether, we believe that this expanded toolkit of available screening technologies will enable an informed selection of the most appropriate technology for CMOD screening, depending on whether speed, spatial information or dynamics is the priority. This will enable exploration of larger biological and chemical space at scale, allowing identification of CMODs with unique mechanisms of action tailored to the specific condensate biology in question.

## Results

### htSMT can report on GSK3 inhibitor modification of N protein dynamics

Single molecule tracking can be used to measure changes in protein-protein and protein-nucleic acid interactions in cells. Because condensate formation is thought to arise through an increase in multivalent interactions, we hypothesized that the pro-condensation effects of GSK3 inhibitors on SARS-CoV-2 N could be measured with htSMT. To demonstrate this, we began by measuring changes in N dynamics in response to the ATP-competitive GSK3 inhibitor LY2090314. We generated stable U2OS cells expressing N fused to the HaloTag domain via a TEV linker (N-Halo) and validated that this stable cell pool can form visible N condensates upon 24h treatment with 10µM LY2090314 with fluorescence microscopy (Supplementary Figure 1A).

We next performed the htSMT assay with the U2OS N-Halo cells in 384-well plates. Cells were treated with DMSO or 10µM of the positive control GSK3 inhibitor LY2090314 for 24h, then stained with Hoechst and two HaloTag ligands (HTLs) – JF549-HTL (“bulk condensate label”) and JFX650-HTL (“sparse SMT label”) for 1h. Cells were washed, re-treated with compound, and then imaged as described previously^43^ (Figure 2A). Bulk condensate labeling was used to visualize organized N protein structures in the cell, including larger, emergent structures such as phase-separated condensates (Supplementary Figure 1B). Within a single field of view (FOV), N protein dynamics were quantified with the sparse JFX650-HTL SMT label by detecting single molecules (Figure 2B) and temporally connecting them into single molecule trajectories (Figure 2C; Supplementary Movies 1A, B). Within FOVs, jump lengths from individual trajectories were aggregated and used to derive downstream screening metrics (Figure 2D, top; Supplementary Table 1). First, we used the median jump length as the most direct measurement of particle motion, independent of the underlying protein behavior. Similarly, we used the particle tracks to determine the average diffusion coefficient (*D̂*) per FOV, a measurement frequently used in SMT analyses which may also be compared to other biophysical techniques like fluorescence correlation spectroscopy (FCS). Finally, we applied the State Array algorithm, a powerful analysis method, to infer the relative frequency of different N mobility states (Figure 2D, bottom; Supplementary Table 1). These three metrics are correlated, but each give increasing insight into the underlying behavior of the protein under study.

**Figure 2:**
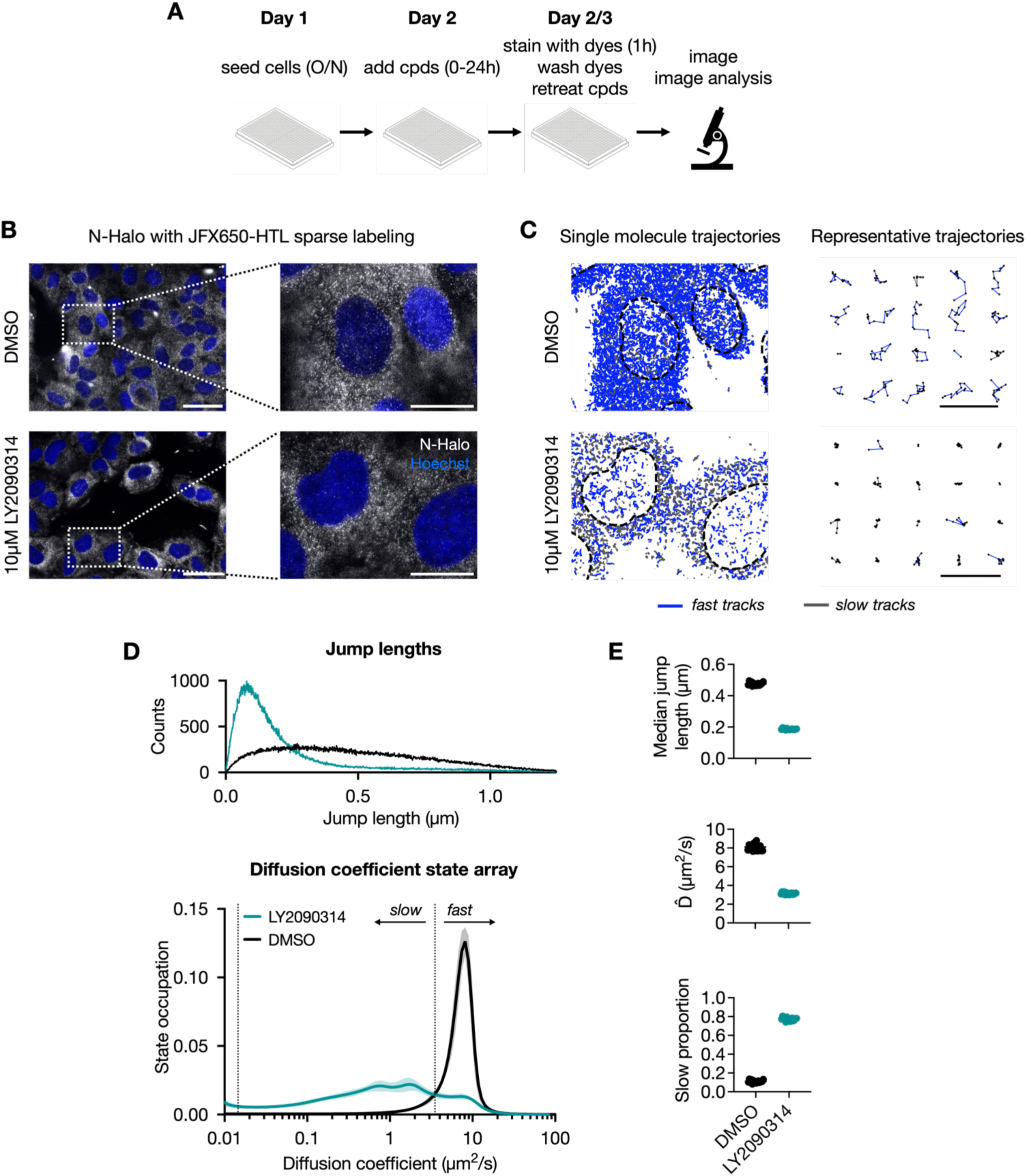
Application of high-throughput single molecule tracking (htSMT) for reporting on GSK3 inhibitor-dependent slowing of N protein dynamics. *A.* Workflow for identifying CMODs that induce N condensation by htSMT. O/N: overnight; cpds: compounds. *B.* Representative FOVs (left) of U2OS N-Halo cells sparsely labeled with JFX650-HTL during SMT image acquisition, upon treatment with DMSO or 10µM LY2090314 for 24h. Each FOV captures 10-40 cells. Four FOVs are captured per well, and for each experiment, anywhere from four to 76 wells of replicates are imaged and analyzed. The insets (right) depict a zoomed-in view illustrating single N-Halo molecules labeled with JFX650-HTL. Full FOV scale bar = 50µm; inset scale bar = 20µm. *C.* Trajectory maps illustrating single N molecule trajectories upon treatment of U2OS N-Halo cells with DMSO or 10µM LY2090314 for 24h. Left: single N molecule trajectories overlayed on Hoechst-delineated nuclei masks (dotted lines). Right: representative trajectories for each treatment condition. Slow-moving (gray) and fast-moving (blue) N molecules are illustrated (see Supplementary Figure 1C and Materials and Methods for definition of slow and fast). Representative trajectories scale bar = 5µm. *D.* Jump length distributions and diffusion coefficient state arrays for characterizing the effect of CMODs on N dynamics. Cells were treated either with DMSO or 10µM LY2090314 for 24h, and then imaged by htSMT. Single N molecule trajectories were used to calculate jump lengths (left). The jump length distribution profiles were then used to infer a diffusion coefficient state array illustrating the probability of various diffusion states within the population (right). The slow and fast proportions (see Supplementary Figure 1C and Materials and Methods for definition) are also represented on the state array, with the dotted lines illustrating the boundaries separating the slow from the fast population. *E.* Dot plot illustrating htSMT metrics – median jump length, *D̂* (average diffusion coefficient), and slow proportion of N molecules treated with DMSO or 10µM LY2090314 for 24h from (D).

Consistent with the expectation that GSK3 inhibition would lead to N self-interaction, upon treatment of cells with 10µM LY2090314 for 24h, there was a visible decrease in the jump lengths of N (Figure 2C). The median jump length from 0.48µm to 0.19µm, and *D̂* from 8.1µm^2^/s to 3.15µm^2^/s (Figure 2E, Supplementary Table 1). Looking at the diffusion coefficient state arrays, we observed a dramatic shift to at least one low mobility state. To further highlight the distinct N mobility states during CMOD treatment, we developed a third metric coined “slow proportion” to describe the abundance of N moving to these slower states (Figure 2D, E). This was done by searching for a pair of diffusion coefficient boundary values which maximally differentiated the two control conditions using Z’ score. (Supplementary Figure 1C; Supplementary Table 2; *Materials and Methods*). After differentiating the two mobility states based on their state distributions, we observed an increase in the slow proportion of N from 0.11 to 0.77 after treatment with LY2090314, consist with the trends in decreasing median jump length and *D̂*(Figure 2E, Supplementary Table 1). In contrast with the large difference in dynamics, we observed only a modest increase in visible N condensate puncta induced by LY2090314 through simultaneous imaging of bulk N labeling via JF549-HTL (Supplementary Figure 1B). These data suggest that htSMT is more sensitive to differences in N self-interaction than bulk N condensation under these specific assay conditions. These changes in dynamics were reproduced in a separate HeLa N-Halo clonal cell line, demonstrating that multiple cell lines are compatible with htSMT (Supplementary Figure 2A-C, Supplementary Table 3). Taken together, these data show that the pro-condensation phenotype of the GSK3 inhibitor LY2090314 was reliably quantified using the htSMT platform.

### Structure-activity relationships (SAR) from htSMT data

We further tested two additional GSK3 inhibitors, CP21R7 (ATP-competitive) and tideglusib (allosteric inhibitor), in the htSMT assay to evaluate any differences in N dynamics compared to LY2090314 treatment. As expected, CP21R7 treatment resulted in slow N diffusion (Figure 3A; Supplementary Table 4), as quantified by a decrease in median jump length, average diffusion coefficient *D̂*, and an increase in the slow proportion of N (Figure 3B; Supplementary Table 4). Conversely, tideglusib treatment did not result in changes to any of the three screening metrics compared to DMSO treatment. These findings are consistent with our previous observation that ATP-competitive, but not allosteric GSK3 inhibitors are N pro-condensation CMODs resulting in reduced N dynamics.

**Figure 3:**
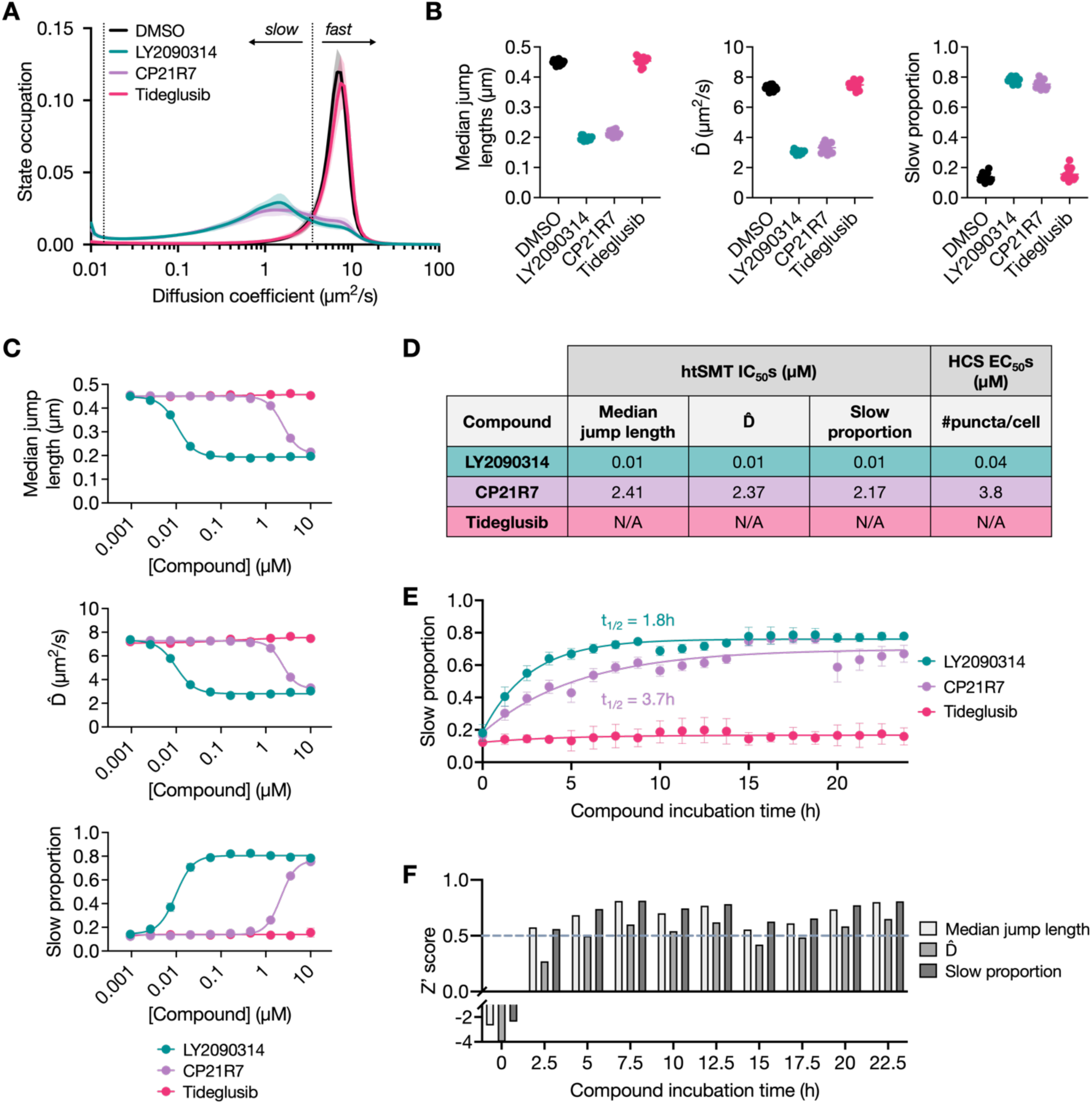
htSMT can sensitively report on changes in SARS-CoV-2 N dynamics in response to pro-condensation GSK3 inhibitors. A. Diffusion coefficient state array illustrating diffusion state probability distributions for a U2OS N-Halo-expressing stable pool treated with DMSO or 10µM LY2030314, CP21R7, or tideglusib for 24h. The slow and fast populations are also represented on the state array, with the dotted lines illustrating the boundary. B. Dot plots indicating jump lengths, *D̂*, and slow proportions for 24h treatment with DMSO or 10µM LY2030314, CP21R7, or tideglusib. C. Dose response curves illustrating jump lengths, *D̂*, and slow proportions for 24h treatment with DMSO, LY2090314, CP21R7 or tideglusib at different concentrations. D. Table comparing IC50/EC50 values for each of the three metrics in comparison to the EC50 values of N condensation observed by previously reported HCS. E. Time course plot illustrating changes in slow proportion of N over time after treatment with DMSO or 10µM LY2090314, CP21R7, or tideglusib. Data from a continuous htSMT assay were binned per hour. F. Time course plot illustrating Z’ scores over time for each of the three metrics. Dotted line indicates the desired Z’ score cut-off of 0.5.

To determine the relative potencies of each GSK3 inhibitor in reducing N dynamics, we assessed them in dose response format and measured N dynamics following a 24h treatment. As earlier, the allosteric inhibitor tideglusib did not affect the jump length, *D̂*, or slow proportion of N (Figure 3C; Supplementary Table 4). For the ATP-competitive inhibitor LY2090314, the EC_50_ values calculated for all three screening metrics (jump length, *D̂*, slow proportion) were the same at 0.01µM (Figure 3C-D; Supplementary Table 4). Similarly, the potency values for the three screening metrics were comparable at 2.41µM, 2.37µM, and 2.17µM for CP21R7 (Figure 3C-D; Supplementary Table 4). Importantly, these recorded EC_50_ values were comparable to previously reported EC_50_ values from puncta counting in HCS for all three compounds, with LY2090314 most potently reducing N protein dynamics (Figure 3D). These results were reproduced at 6h post treatment for all three screening metrics (Supplementary Figure 3A-B; Supplementary Table 5), highlighting the sensitivity of htSMT in resolving condensation dynamics even at time points when conventional HCS approaches would struggle. Altogether, this suggests that monitoring changes in N protein dynamics through htSMT can give insights into N condensation and is a viable alternative readout for monitoring phase separation events starting at early time points.

Next, we conducted a kinetic analysis of changes to htSMT readouts upon GSK3 inhibition where changes in N dynamics were monitored continuously over 24h following treatment with the three CMODs (Figure 3E; Supplementary Table 4). As expected, 10µM tideglusib did not result in changes to N dynamics over the full assay window. In contrast with tideglusib, both 10µM LY2090314 and CP21R7 treatments resulted in the slow proportion of N increasing from less than 20% to a plateau of above 70%. Interestingly, the slow proportion of N increased at a faster rate for LY2090314 treatment (t_1/2_ = 1.8h) as opposed to CP21R7 treatment (t_1/2_ = 3.7h), highlighting its enhanced potency in driving N condensation.

Further, we sought to analyze how htSMT metrics perform in a screening format over time. Using LY2090314 as a positive control, we calculated Z’ scores for jump length, *D̂*, and slow proportion over a continuous period of 24h using bins of 1 hour of collected data. Using a Z’ score threshold of greater than 0.5 as an indicator of a robust screening assay, we found that 2.5h of treatment with 10µM LY2090314 resulted in a difference in N dynamics sufficient for robust screening (Figure 3F; Supplementary Table 4). Notably, Z’ scores were always higher for the raw jump length and the slow proportion metric as compared to *D̂*(Figure 3F). A similar time-course experiment was performed with the traditional HCS assay pipeline with fixation and staining with JF479-HTL to quantify N puncta upon 24h treatment with LY2090314 (Supplementary Figure 4A; Supplementary Table 6). We found that the difference in the number of puncta between the LY2090314-treated cells and DMSO-treated cells is small at all time points throughout the 24h assay. This difference is underscored by the calculated Z’ scores, which did not surpass the Z’ threshold of 0.5 (Supplementary Figure 4B). Altogether, this highlights the improved sensitivity of htSMT in detecting early changes in protein dynamics in comparison to coarser changes in N protein localization and organization.

### Proximity-based condensate biosensors also report on CMOD-induced N condensation

To develop a fast plate-reader assay for characterizing CMODs, we designed proximity-based biosensors that report on protein condensation with a simple plate reader readout, independent of any imaging modality. To generate N condensate biosensors, we fused the large (Lg) and small (Sm) subunits of the split-NanoLuc enzyme, or the NanoLuc donor and HaloTag acceptor domains to the C terminus of the N protein (Figure 4A). For this high-throughput assay, HEK293T cells were first seeded in large dishes, transfected with reporter constructs and replated into 384-well plates the following day. Cells were then treated with the three GSK3 inhibitors for 24h before performing the plate reader assay (Figure 4B).

**Figure 4:**
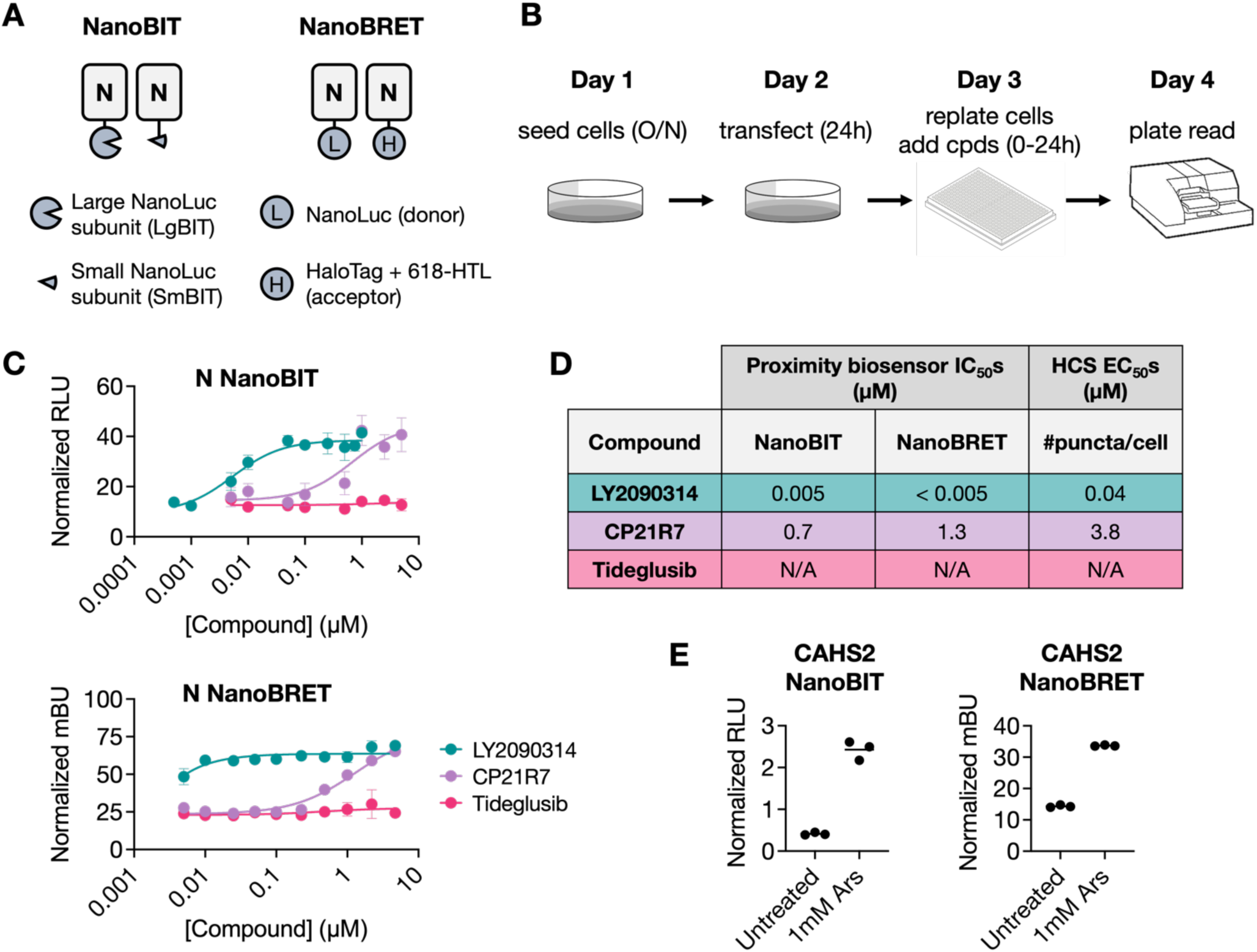
Application of proximity-based condensate sensors for identification of CMODs. A. Schematic illustrating the proximity-based NanoBIT and NanoBRET N condensate biosensors. Both systems provide a readout when N molecules are in nanometer-scale proximity, such as in the condensed phase as opposed to the dilute phase. For the NanoBIT system, the NanoLuc enzyme is only functional when the large and small NanoLuc subunits interact in proximity. For the NanoBRET system, the acceptor 618-HTL fluorophore can only be excited when in proximity to the donor NanoLuc. B. Workflow for identifying CMODs that induce N condensation by proximity-based condensate sensors. O/N: overnight; cpds: compounds. C. Dose response curves illustrating NanoBIT and NanoBRET signals for treatment of N reporters with DMSO, LY2090314, CP21R7, or tideglusib at different concentrations. D. Table representing EC50 values of N condensation for both the NanoBIT and NanoBRET systems in comparison to the EC50 values observed by previously reported HCS. E. Dot plots indicating NanoBIT and NanoBRET signals for treatment of CAHS2 reporters with 1mM sodium arsenite (Ars).

We performed dose response analyses for the NanoBIT and NanoBRET assays. Both LY2090314 and CP21R7 increased the NanoBIT and NanoBRET signals in a dose-dependent manner, whereas tideglusib left the signals unchanged (Figure 4C; Supplementary Table 7). Importantly, the EC_50_ values recorded for LY2090314 and CP21R7 were 0.006µM and 0.7µM for the NanoBIT reporter, and <0.005µM and 1.3µM for the NanoBRET reporter respectively (Figure 4D). These trends are consistent with the observed potencies of these CMODs based on previous HCS assays and htSMT, where LY2090314 most potently induced N condensation. We also validated the formation of N puncta in the HEK293T cells expressing the reporter constructs by immunofluorescence and confirmed the formation of N puncta upon treatment with LY2090314 and CP21R7 but not tideglusib (Supplementary Figure 5A).

To demonstrate the applicability of these proximity-based condensate sensors beyond the SARS-CoV-2 N protein, we tested the two systems with the tardigrade cytosolic abundant heat-soluble (CAHS) protein CAHS2. We found that treatment of cells with sodium arsenite, an inducer of oxidative stress and stress granule formation in human cells, resulted in an increase in CAHS2 NanoBIT and NanoBRET signals (Figure 4E; Supplementary Table 7). Formation of intracellular condensates was similarly confirmed by immunofluorescence (Supplementary Figure 5B), pointing towards oxidative stress as a distinct form of cellular stress that can induce morphological changes in CAHS2 organization. Taken together, these data suggest that the NanoBIT and NanoBRET proximity-based assays can readout changes in protein condensation states. As such, these assays are suitable reporters for modulation of protein condensation beyond SARS-CoV-2 N protein and can be applied for high-throughput screening to identify novel protein CMODs for other targets.

## Discussion

To further advance technologies available for CMOD discovery, we developed two alternative assays, high-throughput single molecule tracking (htSMT) and proximity-based condensate biosensors, for identification of phase separation modulators in a small molecule screening context. CMODs have distinct mechanisms to alter condensate biology, broadly either by altering actual condensation states, such as through changing post-translational modifications or expression levels of condensation-associated proteins, or by altering the biochemistry within condensates^25^. Given the considerable potential of condensate modulation by small molecules, there have been numerous efforts to identify CMODs through large-scale chemical screens. However, these screens are traditionally performed with phenotypic HCS, which suffer from several limitations, including lack of information on condensate dynamics, poor throughput and resolution, and inaccuracies in image analysis pipelines. Most of these limitations arise from the need to identify and quantify condensate puncta as the screening output.

Our two alternative assay modalities to identify CMODs have complementary advantages for high-throughput screening in condensate biology especially since they report on molecular level interactions instead of emergent condensation phenotypes (Supplementary Table 8). Briefly, HCS can be performed on cell lines without cell line engineering with immunofluorescence imaging, allowing it to be rapidly adaptable to studying native condensation biology. On the other hand, htSMT can be useful for reporting on both self-interactions as well as interactions with other scaffold or client proteins (known or unknown), since both types of interactions would result in the slowing of protein motion. Furthermore, capturing single molecule dynamics can be applied to studying CMOD perturbations that do not change the emergent condensation phenotype. The proximity-based biosensors can be used to rapidly probe self-interactions with a simple readout, which is useful in investigations where scale is favored over information content. Altogether, this provides an expanded toolkit for dissecting complex molecular interactions in condensate biology.

SMT has previously been harnessed to illuminate the role of protein dynamics in condensate biology^44–46^, particularly in the fields of transcriptional condensates^47^ and synaptic vesicle organization and function^48^. We hypothesized that measuring changes in dynamics of condensing proteins can inform on the condensate modulatory activity of small molecules and may serve as a useful screening modality for CMODs and other small molecules. Our htSMT assay with the SARS-CoV-2 N protein highlighted not only the scalability of SMT and super resolution microscopy, but also that changes in N protein dynamics can be robustly correlated with emergent condensation phenotypes. We and others previously showed that GSK3 inhibition by the various CMODs investigated led to decreased phosphorylation of N and hence condensation^32, 49, 50^. As such, our htSMT results are consistent with either (a) dephosphorylation-induced oligomerization of N en route to phase separation (i.e. larger oligomer hydrodynamic radius), and/or (b) phase separation itself (i.e. N existing in a densely packed milieu) giving rise to a decrease in N dynamics. Of note, the decrease in dynamics observed in our assays were independent of any observable changes in N condensation through traditional fluorescence imaging.

Our htSMT assays highlight several advantages over traditional HCS assays. Firstly, htSMT revealed the rate at which CMODs induce observable changes in protein dynamics, which can be used to infer any mechanistic differences between CMODs. For example, we observed that LY2090314 resulted in a faster increase in the slow proportion of N than CP21R7, consistent with the possibility that LY2090314 may exhibit a higher binding affinity to GSK3. Secondly, htSMT can enable the study of smaller condensate formations and protein state changes that cannot typically be observed by fluorescence microscopy. Finally, we observed that robust Z’ scores can be achieved as early as 2.5h post-CMOD addition. This allows for htSMT screening to be performed in more sensitive systems, for example in cell lines where compound treatment for a prolonged time may result in cell toxicity. There are also several additional advantages of htSMT over traditional HCS that were not explored in our study. Firstly, hSMT can identify CMODs that perturb condensate dynamics without altering the number, size, or morphology of the condensates. These CMODs would typically not be identified by image analysis in traditional HCS. Secondly, unique distribution profiles in a diffusion coefficient state array can indicate distinct mechanisms of action of CMODs (for example, a shift in the distribution peak compared to a flattening of the peak). Altogether, our htSMT assay provides robust information on N condensation states and good Z’ scores, making it suitable for further condensate biology characterization and CMOD high-throughput screening.

High-throughput plate reader assays are valuable for applications where a larger pilot screen is valued over the amount of information collected from the initial screen. With our proximity-based biosensors, the screening readout can be completed in <5 min as compared to over an hour long with HCS, per 384-well plate. Since the readout is imaging-agnostic, the assay reports on changes in molecular-scale interactions without being subjected to inaccuracies in image analysis pipelines. We also validated that this proximity-based sensor can read out bulk condensation states even when the condensing protein exists as a dimer. From our results, we still observed a dose-dependent increase in signal for both the NanoBIT and NanoBRET systems for N in the presence of the ATP-competitive GSK3 inhibitors, despite the baseline level of proximity-induced signal due to N dimerization. This could be because not all N molecules are dimerized in the untreated state, and specifically in case of the NanoBRET system, the induced proximity with condensation can lead to a non-stoichiometric excitation of more than one acceptor fluorophore per donor luminescence. This also underscores the importance of optimizing both the overall and the stoichiometric expression level of the condensate sensor pairs in cells. Too high of an overall expression level (i.e. with the use of a strong promoter), in some cases, may lead to a lower dynamic range due to higher background signal. Changing the ratio of each of the condensate sensors to their partners can also affect the amount of background signal by modulating the number of functional dimers in the absence of CMODs. As such, assay optimization is critical for application of proximity-based condensate sensors to new target condensing proteins to determine the best expression conditions where the dynamic range between the positive and negative controls is maximized.

To demonstrate that the NanoBIT and NanoBRET sensors can be applied to condensing proteins beyond SARS-CoV-2 N, we showed that several other HCoV N proteins and the tardigrade protein CAHS2 can respond to known and novel CMODs. CAHS2 is part of a larger family of CAHS proteins known to protect tardigrades against stresses such as desiccation by forming intracellular protective fibrils and gels^51^. We discovered that sodium arsenite-induced stress triggered the phase separation of CAHS2 and hence an increase in the NanoBIT and NanoBRET signal. This raises the possibility that CAHS2 condensates may function as the tardigrade equivalent of stress granules under heavy metal or oxidative stress conditions, thereby also illustrating the utility of this system for rapidly interrogating novel condensate biology.

In summary, our work advances the field of high-throughput screening for CMODs by introducing and validating two innovative technologies – htSMT and proximity-based condensate biosensors – that overcome some of the limitations of traditional HCS methods by reporting on molecular interactions instead of the condensation phenotype. We believe that this versatile toolkit of complementary technologies will be instrumental in designing and optimizing CMOD screening assays tailored to specific condensate biology. This lays the foundation for future studies to explore the potential of condensate-targeted therapies in combating a wide range of diseases, from neurodegeneration to cancer and beyond.

## Materials and methods

### Cell lines and cell culture

HEK293T/17 (CRL-11268), A549 (CCL-185), U2OS (HTB-96) and HeLa (CCL-2) cells were purchased from ATCC. HEK293T/17, U2OS and HeLa cells were maintained in Dulbecco’s modified Eagle’s medium (DMEM ATCC 30-2002 or Gibco 10566016) supplemented with 10% (v/v) fetal bovine serum (FBS; Gibco 10438026 or Corning 35-015-CV), penicillin-streptomycin (Gibco 15140122 or 10378016) and normocin (Invivogen ant-nr-1). A549 cells were maintained in F-12K medium (ATCC 30-2004) supplemented with 10% (v/v) FBS, penicillin-streptomycin and normocin. All stable cell pools or clonal cell lines were maintained in full DMEM culture medium supplemented with 1.5µg/mL puromycin (Gibco A1113803). Cells were maintained at 37°C and 5% CO_2_ in a humidified environment. Cells were subcultured twice a week by DPBS washing (Gibco 14190250) followed by trypsinization (Corning MT25053CI) from 90% to 20% confluence.

### Plasmid construct generation

The pHAGE lentiviral plasmid encoding SARS-CoV-2 N-TEV-Halo-V5 (for generation of htSMT cell lines), SARS-CoV-2 N-Nluc and N-Halo (for NanoBRET) were synthesized by Twist, generating rt256a, rt162a and rt163a respectively. For NanoBIT constructs, the pHAGE lentiviral plasmid encoding SARS-CoV-2 N-LgBIT and N-SmBIT were generated by NEBuilder HiFi DNA Assembly (New England Biolabs E2621S). Oligonucleotide primers were obtained from Azenta Life Sciences. The SARS-CoV-2 N sequence and the NanoBIT vector backbones pBIT1.1-C [TK/LgBIT] and pBIT2.1-C [TK/SmBIT] were amplified by PCR (New England Biolabs M0492S) and cloned by NEBuilder HiFi DNA Assembly, generating N-LgBIT and N-SmBIT regulated by the HSVTK promoter. To standardize backbone contexts, the entire HSVTK-N-LgBIT, HSVTK-N-SmBIT and negative control HSVTK-Halo-SmBIT fragments were amplified and cloned into the pHAGE backbone, generating rt188a, rt189a and rt190a respectively. This replaced the original CMV promoter in the pHAGE backbone for optimal expression levels of these constructs for our assays. The puromycin selection marker from rt162a and rt188a was replaced with a blasticidin selection marker amplified from pcDNA™6/TR (Invitrogen V102520), generating rt192a and rt191a respectively. To generate NanoBIT and NanoBRET constructs for SARS-CoV N, HCoV-OC43 N, MERS-CoV N and CAHS2, the genes were amplified from plasmids from previous studies^32, 52^ and used to replace the SARS-CoV-2 N gene in rt189a/rt191a (NanoBIT pair) and rt163a/rt192a (NanoBRET pair) by cloning. Plasmid construct details can be found in Supplementary Table 9.

### Stable cell line generation

Stable U2OS and HeLa cell lines expressing SARS-CoV-2 N-Halo were generated by lentivirus transduction as previously described^32^ using the rt256a construct. After selection recovery and expansion, cells were sparsely seeded in 15cm tissue culture dishes (50-100 cells/dish) and clonal cell lines were harvested with cloning cylinders (Millipore Sigma C1059). Both U2OS and HeLa stable pools and clonal cell lines were expanded and stored in liquid nitrogen.

### Fluorescence imaging-based assays and image analysis

Where referenced, fluorescence imaging-based HCS assay results obtained with A549 cells stably expressing SARS-CoV-2 N-EGFP were obtained from our previous study^32^. For kinetic imaging-based assays performed with the U2OS N-Halo stable cell pool, cells were plated on multiple 384-well PhenoPlates (Revvity 6057302) at 4,000 cells/well and treated with DMSO or 10µM LY2090314. At specified time points post-compound addition, each plate was fixed with 4% (w/v) formaldehyde in PBS (Thermo Fisher Scientific 28908) with 1µg/mL Hoechst 33342 (Invitrogen H3570) and 0.5µM JF479-HTL (Janelia Research Campus) for 30min at room temperature before washing 3x with PBS. Plates were imaged with MetaXpress (version 6.7) on a Molecular Devices ImageXpress Micro Confocal Laser (IXM-C LZR) microscope equipped with a spinning disk confocal scanner, a high-quantum efficiency sCMOS detection camera and a Lumencor Celesta light engine. Plates were imaged with a 40x/0.95 air objective lens (Nikon) at a pixel size of 0.16µm/pixel. Images were collected with a 477nm laser, 488nm dichroic filter and 520/25nm emission filter Hoechst staining was imaged with a 405nm laser, 421nm dichroic filter and 452/45nm emission filter. Images were processed in FIJI (NIH).

### htSMT assay

U2OS N-Halo stable pools and HeLa N-Halo clonal cell lines were thawed from assay-ready media (90% FBS, 10% DMSO) into full DMEM culture media. Cells were plated on 384-well CellVis plates at 6,000 cells/well for 6h treatment experiments and at 4,000 cells/well for 24h treatment experiments.

Cells were treated either in an 11-point 3-fold dose response with compounds from 10µM to 0.001µM (for dose response experiments) or a single 10µM dose of compounds (for kinetic experiments), that was transferred from Echo Qualified 384-Well Low Dead Volume Source Microplate (Beckman Coulter 0018544) and incubated at indicated time points at 37°C. Cells were then stained with 100nM Hoechst 33342 (Invitrogen H3570), 200pM JF549-HTL dye (Promega GA1110) for bulk labelling, where specified, and 1.56pM JFX650-HTL dye for SMT (Promega HT1070). Cells were then incubated for 1h at 37°C before being washed using the EL406 microplate washer (Aligent BioTek) into imaging media composed of FluoroBrite™ DMEM media (Gibco A189670) supplemented with GlutaMAX (Gibco 35050079) and the same serum and antibiotics as culture media.

### htSMT imaging

All images were acquired with Oblique line scanning (OLS) microscopy as previously described^43^. Briefly, cells were imaged at 100 Hz (0.01 second frame intervals) for 150 frames of SMT with 642nm excitation for JFX650 tracking. Masking for Hoechst (nuclear staining) was imaged with 405nm excitation and was collected as a single frame for downstream nuclear segmentation. Bulk N labeled with JF549 was imaged with 561nm excitation and was collected as a single frame for qualitative assessment of N-protein condensate puncta formation, where indicated. For kinetic experiments, multiple plates were set up as described above and imaged over the course of 24h.

### htSMT image analysis

Particle detection and localization methods used are identical to those previously described^43^. Localized particles were linked with a proprietary variational Bayesian, non-sampling-based inference routine which simultaneously optimizes particle links and estimates individual trajectory diffusion coefficients. To limit linking error, an upper bound of 1.25µm jump lengths and 2 frame gaps were permitted. Localized detections, or spots, were automatically filtered based on shape and intensity characteristics to limit the number of false detections due to camera noise, fluorescent cell debris, free dye, or other non-single-emitter signals. Individual fields of view were also filtered based on the number of segmented cells to discard movies with sparse cells or focus problems.

“Dye-like” trajectories were used as inputs for resolving a spectrum of diffusive states using the State Array algorithm on a per-FOV basis^53^. “Regular Brownian Motion with Localization Error” (RBME) State Arrays were run as previously described^43^ with the exception that track aggregation was done per FOV rather than per well. The minimum and maximum diffusion coefficient bins used for State Arrays were 100 evenly spaced log_10_ bins between 0.01 and 100µm^2^/s, respectively. For U2OS cells, an additional filter of 50,000 tracks per FOV was imposed for more stringent quality control.

The “slow proportion” metric was derived with kinetic experiment data at 14h post-compound treatment because the diffusion coefficient state arrays visually showed at least two distinct diffusive states. These data included 76 negative control (DMSO) wells and 76 positive control (10µM LY2090314) wells with 4 FOVs per well. To derive a diffusion coefficient boundary for “slow proportion”, an exhaustive search of variable diffusion coefficient floors and ceilings was applied to select diffusion coefficient states within the designated window. Then, Z’ scores were calculated using the integrated proportion of diffusion coefficient states within the selected window (Supplementary Figure 1B for U2OS N-Halo; Supplementary Figure 2A for HeLa N-Halo). The maximum-minimum diffusion coefficient pair that led to the highest Z’ score per cell line was selected to define the boundary separating the slow and fast population in the respective cell line.

### Proximity-based assays

HEK293T/17 cells were seeded in 6-well plates at 800,000 cells/well and transfected with 20µg of each plasmid of a pair (40µg total per well). After 24h, cells were trypsinized and replated into 96-well (Corning 3610) or 384-well (Corning 3765) plates at 30,000 cells/well or 6,000 cells/well respectively. On the same day (for NanoBRET) or on the next day (for NanoBIT), cells were treated with compounds in dose response with the HP D300e digital dispenser for 24h. For the NanoBIT assay, the Nano-Glo® live cell assay substrate (Promega N2011) was added to cells according to manufacturer’s protocols and luminescence readings were recorded with the BioTek Synergy H1 plate reader using with an integration time of 0.5s and a gain of 200. It was noted that using half the volume of the substrate did not result in any significant difference in dose response EC50s or dynamic ranges, therefore optimization of substrate concentration/volume is possible. For the NanoBRET assay, the HaloTag-618 ligand (Promega N1662) was added to cells at a final concentration of 100nM before replating. The NanoBRET® Nano-Glo® substrate was added to cells according to manufacturer’s protocols and both the donor (460nm) and acceptor (618nm) emissions were measured with the NanoBRET Blue 460/80nm and BRET Deep Red 647 647/75nm filters respectively and the LUM/D585 mirror on a PerkinElmer EnVision plate reader. All experiments were performed in duplicates or triplicates.

### Proximity-based assays data analysis

For the NanoBIT assays, the raw relative luminescence units (RLU) collected for each well (“sample”) does not consider total cell count in the well, or cell toxicity. An additional transfection condition (“negative”) was performed where LgBIT constructs were co-transfected with a SmBIT construct encoding a non-interacting protein partner, HaloTag-SmBIT. The sample RLU from each well was normalized to the mean RLU of the negative wells to account for cell toxicity (“normalized RLU”). Additionally, the negative condition allows for sequestration of only the LgBIT components upon treatment with a pro-condensation CMOD, resulting in a lower baseline negative RLU and a larger normalized RLU, thereby increasing the dynamic range of the assay. For the NanoBRET assays, the donor Nluc emission (460nm) can be used as a readout for cell toxicity. The NanoBRET signal in milliBRET units (mBU) was calculated by taking a ratio of the donor Nluc emission (460nm) to the acceptor HaloTag-618 ligand emission (618nm), x 1000 (“normalized mBU”).

### Fluorescence microscopy of NanoBIT biosensor-expressing cells

For qualitative fluorescence imaging of NanoBIT sensor-expressing cells, cells were fixed with 4% (w/v) formaldehyde (Thermo Scientific 28906) in PBS for 30min immediately after plate reader readout. After fixation, plates were washed thrice with PBS and blocked in PBS + 0.1% (v/v) Triton X-100 (Sigma-Aldrich X100) + 2% normal goat serum (Abcam ab7481) at room temperature for 1h. Plates were then incubated overnight at 4°C with primary antibodies (mouse anti-LgBIT antibody at 1:750 dilution [Promega N7100]; mouse anti-SARS-CoV-2 N antibody diluted 1:1,000 [Cell Signaling Technology 33717]) diluted in PBS + 0.1% (v/v) Triton X-100 + 0.5% normal goat serum. Plates were washed thrice with PBS before incubation with secondary antibodies (goat anti-mouse IgG (H+L) Cross-Adsorbed Secondary Antibody, Alexa Fluor™ 568 [Invitrogen A-11004] diluted 1:1,000 or goat-anti-mouse IgG (H+L) Highly Cross-Adsorbed Secondary Antibody, Alexa Fluor™ Plus 488 [Invitrogen A32723]) and 1µg/mL Hoechst 33342 in the same dilution buffer for 1h at room temperature. Plates were washed again thrice with PBS before imaging with MetaXpress (version 6.7) on the Molecular Devices IXM-C LZR as described earlier.

## Acknowledgements

This work was sponsored by a research alliance with AbbVie, Inc. Rui Tong Quek was supported by the Agency for Science, Technology and Research NSS (Ph.D.) predoctoral fellowship. The htSMT portions of this work were made possible through participation in the Technology Access Program at Eikon Therapeutics, Inc. The authors thank the many current and former Eikon employees that have enabled this work and provided thoughtful feedback on the data. The content and conclusions included in this manuscript are solely those of the authors. The authors thank the ICCB-Longwood Screening Facility at HMS for access to high-throughput screening equipment.

## Author contributions

R.T.Q, designed constructs and generated cell lines for htSMT and proximity-based condensate biosensors. C.L.Y, C.R.S, P.J.E. performed the htSMT assays and analysis. R.T.Q., K.S.H. and S. L. performed proximity-based biosensor experiments and performed data analyses. R.T.Q. wrote the manuscript. P.J.E., D.T.M, T.J.M. and P.A.S. managed the team, assisted with interpretation of results, and provided guidance on manuscript writing. R.T.Q., C.L.Y, C.R.S, P.J.E. and D.T.M. participated in the study design and execution of htSMT experiments, including data interpretation and manuscript review and approval. All authors provided critical feedback on the manuscript.

