## Supplementary Figures for "Comparative evaluation of cell-based assay technologies for scoring drug-induced condensation of SARS-CoV-2 nucleocapsid protein"

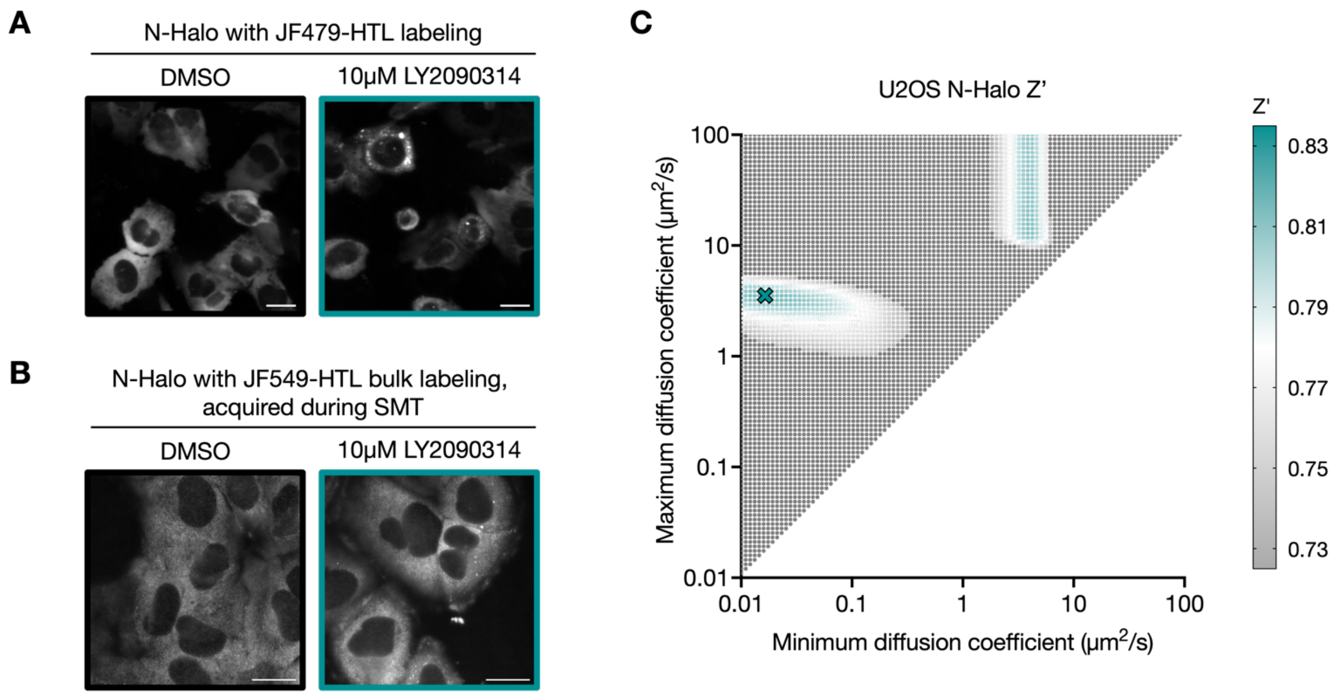


***Supplementary Figure 1: htSMT assay validation and description of slow and fast proportions of N molecules.***

A. *Fluorescence images illustrating visible N-Halo puncta upon 24h treatment of U2OS N-Halo stable cells with 10μM LY2090314. Cells were fixed and stained with 0.5μM JF479-HTL for visualization of N condensates. Scale bar = 20μm.*

B. *Fluorescence images illustrating bulk condensate labeling with JF549-HTL for identification of phase separated N condensates during htSMT. Images represent cells treated with 10μM LY2090314 for 24h.*

C. *Matrix for determining slow proportion boundaries in U2OS N-Halo cells. The matrix represents Z' scores for each combination of a minimum and maximum diffusion coefficient value from the diffusion coefficient state array in Figure 2E. The proportion of molecules falling within the inclusive bounds of each pair of minimum and maximum value combinations was used to calculate the Z' score using DMSO treatment as the negative control and LY2090314 as the positive control. Under these assay conditions, defining the slow proportion as the proportion of molecules falling within the inclusive 0.016μm$^2$/s and 3.51μm$^2$/s diffusion coefficient boundaries led to the highest Z' score values. This minimum-maximum diffusion coefficient pair representing the bounds of the slow proportion is denoted with a teal cross, where the Z' score is 0.82.*

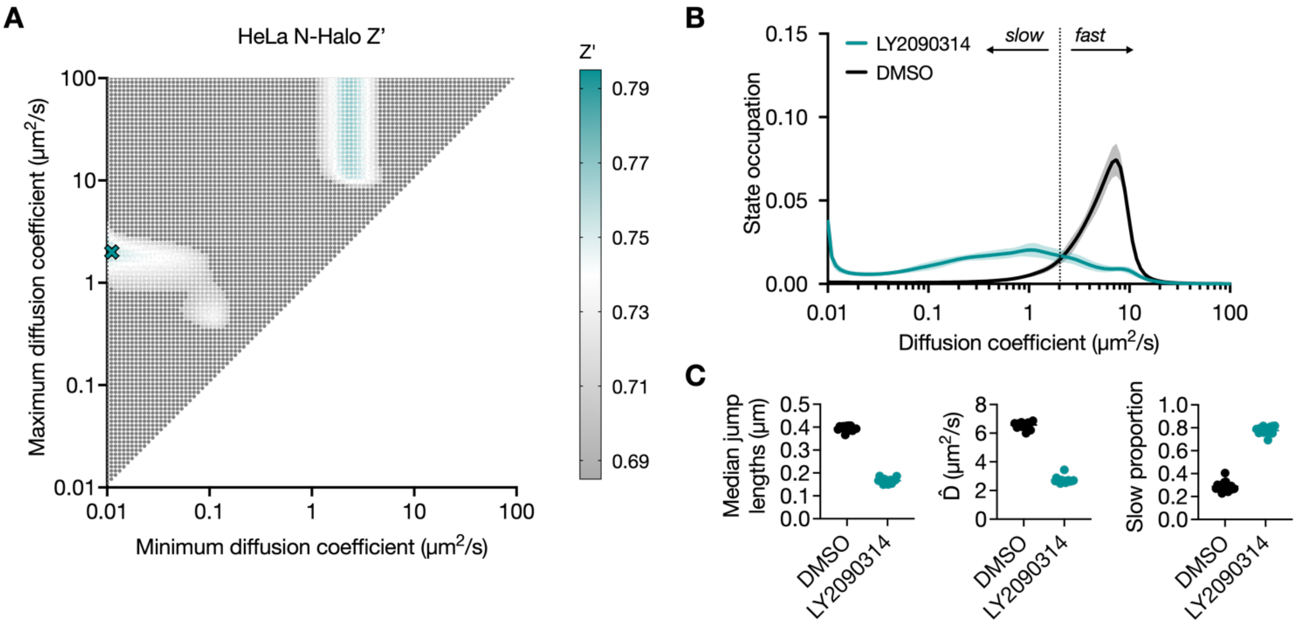


***Supplementary Figure 2: Validation of htSMT in HeLa N-Halo cell line.***

A. *Matrix for determining slow proportion boundaries in HeLa N-Halo cells. The matrix represents calculated Z' scores for each combination of a minimum and maximum diffusion coefficient value from the diffusion coefficient state array in (B). The proportion of molecules falling within the inclusive bounds of each pair of minimum and maximum value combinations was used to calculate the Z' score, with DMSO treatment as the negative control and LY2090314 as the positive control. Under these assay conditions, defining the slow proportion as the proportion of molecules falling within the 0.01μm$^2$/s and 2.01μm$^2$/s diffusion coefficient boundaries led to the highest Z' score values. This minimum-maximum diffusion coefficient pair representing the bounds of the slow proportion is denoted with a teal cross, where the Z' score is 0.77.*
B. *Diffusion coefficient state array illustrating diffusion state probability distributions for a HeLa N-Halo cells treated with DMSO or 10μM LY2030314 for 24h. The slow and fast proportions are also represented on the state array, with the dotted line illustrating the boundary separating the slow from the fast population.*
C. *Dot plot representing htSMT metrics collected for DMSO and 10μM LY2090314 treatment of the HeLa N-Halo cell line in (B).*

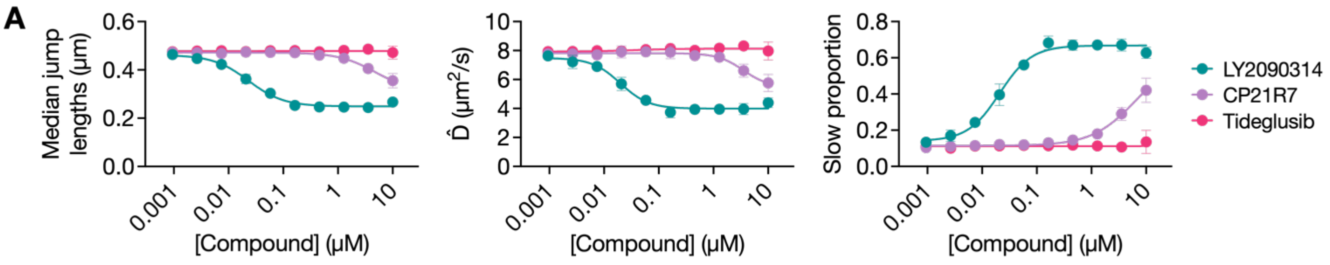


**B**

| | htSMT $IC_{50}$s (μM) | | | HCS $EC_{50}$s (μM) |
|---|---|---|---|---|
| **Compound** | **Median jump length** | **D̂** | **Slow proportion** | **#puncta/cell** |
| **LY2090314** | 0.02 | 0.02 | 0.02 | 0.04 |
| **CP21R7** | 4.8 | 3.5 | 6.6 | 3.8 |
| **Tideglusib** | N/A | N/A | N/A | N/A |

***Supplementary Figure 3: htSMT assay with 6h compound treatment.***

A. *Dose response curves illustrating jump lengths, $\hat{D}$, and slow proportions for 6h treatment of U2OS N-Halo cells with DMSO, LY2090314, CP21R7 or tideglusib at different concentrations.*

B. *Table comparing $IC_{50}$/$EC_{50}$ values for each of the three metrics in comparison to the $EC_{50}$ values of N condensation observed by previously reported HCS.*

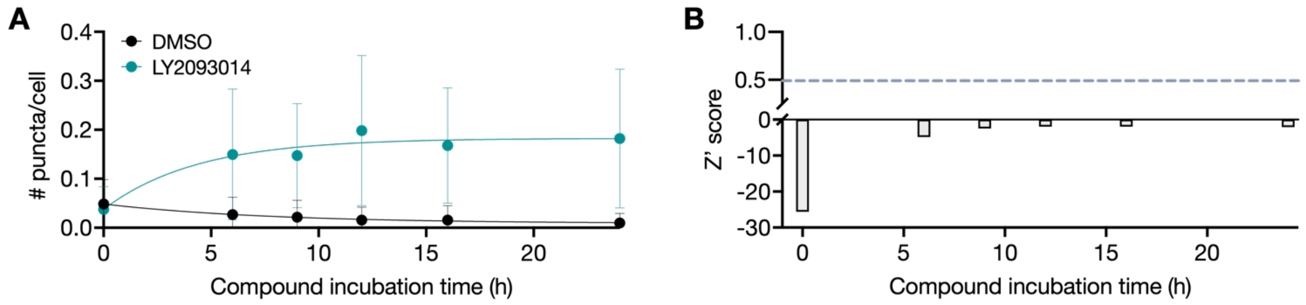


***Supplementary Figure 4: Kinetic imaging-based assay with U2OS N-Halo cells.***

A. *Time course plot illustrating changes in N puncta per cell over time after treatment of U2OS N-Halo cells with DMSO or 10μM LY2090314, as assessed by traditional fluorescence microscopy.*

B. *Time course plot illustrating Z' scores post-treatment with compounds. Dotted line indicates an acceptable Z' score of 0.5.*

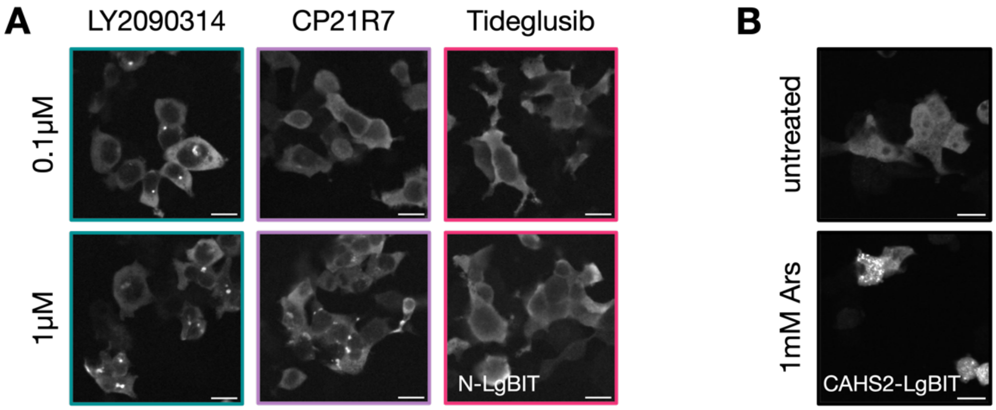


***Supplementary Figure 5: Immunofluorescence imaging validation of phase separation of the various NanoBIT and NanoBRET reporters.***

*A. Fluorescence images demonstrating N protein condensation upon treatment with LY2090314 and CP21R7 but not tideglusib. Cells expressing the NanoBIT reporters were fixed and stained with anti-N antibody for imaging. Scale bar = 20μm.*

*B. Fluorescence images demonstrating CAHS2 protein condensation upon treatment with 1mM sodium arsenite (Ars). Cells expressing the NanoBIT reporters were fixed and stained with anti-large NLuc subunit antibody for imaging. Scale bar = 20μm.*

***Supplementary Movies 1: Representative movies for SMT of U2OS N-Halo cells upon treatment with 10μM GSK3 inhibitors.***

A. *Representative movie for SMT of U2OS N-Halo cells upon treatment with DMSO for 24h. Movie is at 3 frames per second; scale bar = 50μm.*
B. *Representative movie for SMT of U2OS N-Halo cells upon treatment with 10μM LY2090314 for 24h. Movie is at 3 frames per second; scale bar = 50μm.*
C. *Representative movie for SMT of U2OS N-Halo cells upon treatment with 10μM CP21R7 for 24h. Movie is at 3 frames per second; scale bar = 50μm.*
D. *Representative movie for SMT of U2OS N-Halo cells upon treatment with 10μM tideglusib for 24h. Movie is at 3 frames per second; scale bar = 50μm.*

### SUPPLEMENTARY TABLES

Supplementary Table 8: Table of comparison between htSMT, HCS and proximity-based condensate biosensor CMOD screening technologies

| Metric | htSMT | HCS | Proximity-based condensate biosensors | |
|---|---|---|---|---|
| | | | NanoBIT | NanoBRET |
| **Assay development, optimization** | | | | |
| **Fluorophore/tag compatibility** | Requires tagging of POI to HaloTag | Can be performed without tagging POI if antibody is available | Requires tagging of POI to reporter domains | |
| **Expression construct** | Single construct, POI fused to HaloTag | • None required, endogenous expression if antibody available<br>• Single plasmid, POI fused to fluorophore or tag of choice | • Two plasmids, POI-SmBIT/LgBIT<br>• Single plasmid, POI-SmBIT/LgBIT expressed from same construct | • Two plasmids, POI-Nluc/HaloTag<br>• Single plasmid, POI-Nluc/HaloTag expressed from same construct |
| **Construct optimization** | Tags fused to either N or C terminus of POI, where possible test to ensure condensation phenotype is not affected | | | |
| **Stable vs transient expression** | Monoclonal preferred, stable pool acceptable | Stable and transient expression acceptable | Stable and transient expression acceptable | |
| **Size of transcriptional condensates** | Can study small condensates | Can study condensates above diffraction limit | Can study small condensates | |
| **Readout optimization** | SMT (and bulk labeling) dye concentration | Antibody staining | Nluc substrate concentration | Nluc substrate concentration, HaloTag-618 dye concentration |
| **Expression level** | Condensation highly dependent on expression level of POI, must optimize expression level for every set up to ensure good dynamic range between positive and negative controls | | | |
| **Screening assay** | | | | |
| **Major costs incurred to run assay (approx. per 384-well plate)** | SMT JF-HTL dye, bulk labeling JF-HTL dye | Antibody for staining (where required), dyes for tags (where required) | Luciferase substrate (Promega), cell viability assay | HaloTag-618 dye, luciferase substrate |
| **Acquisition time (approx. per 384-well plate)** | ~1h for image/movie acquisition | ~1h for image acquisition | ~5 min for plate reader readout | |
| **Post-acquisition time (approx. per 384-well plate)** | None | ~1h-overnight for antibody/DNA dye staining | None | |

| | | | | |
|---|---|---|---|---|
| **Compound treatment** | Twice (re-treatment of compounds after JF-HTL dye washout, before imaging) | Usually once (unless HaloTag or SNAP-tag fusion proteins used where dye washout is necessary) | Once | |
| **Fixed/live cells (i.e., can reacquire plate?)** | Live, cannot reacquire plate | Usually fixed, usually can reacquire plate | Live, cannot reacquire plate | |
| **Equipment needed for assay readout** | Eikon OLS microscope | High content fluorescence microscope (e.g. Molecular Devices IXM-C, Zeiss Airyscan, PerkinElmer Operetta CLS, etc) | Monochromator- or filter-based plate reader capable of detecting luminescence | Monochromator-based plate reader capable of quantifying luminescence and fluorescence |
| **Potential source(s) of false positives/negatives** | Autofluorescent artifacts | Autofluorescent artifacts | Luciferase inhibitors | |
| **Information collected and post-processing** | | | | |
| **Primary readout data** | Fluorescence image movies | Fluorescence images | Plate reader luminescence values | Plate reader luminescence values, fluorescence values |
| **Image analysis** | Yes, single molecule identification and trajectory connection | Yes, puncta identification and quantification | None | |
| **User involvement in developing data processing methodology** | Minimal, Eikon-developed image analysis pipelines are largely automated | User-developed puncta-counting pipelines (e.g. Ilastik, MetaXpress, ImageJ, CellProfiler, NIS-Elements etc) | Minimal, simple arithmetic calculations | |
| **Time for data processing primary readout data to processed data (approx. per 384-well plate)** | ~1h | ~5-30 min | ~2 min | |
| **Processed metric for condensation state** | Single molecule diffusion coefficients, slow proportion | Number/size/morphology of puncta | Normalized relative luminescence units (RLU) | Normalized milliBRET units (mBU) |
| **Multiparametric data** — **Per well data** | Cell viability (nuclei count) | Cell viability (nuclei count) | None (additional cell viability assay required, or additional set of compound wells for "negative" screening, see *Materials and Methods*) | Cell viability (donor luminescence values) |
| **Multiparametric data** — **Per cell data** | Yes, cell morphology, POI subcellular distribution | Yes, cell morphology, POI subcellular distribution | None | |

| | | | | |
|---|---|---|---|---|
| | **Per condensate data** | Yes, puncta number, puncta size, puncta morphology, puncta subcellular distribution, POI dynamics (jump lengths, diffusion coefficients, slow proportion) within puncta | Yes, puncta number, puncta size, puncta morphology, puncta subcellular distribution | None |
| | **Per molecule data** | Yes, jump lengths, diffusion coefficients, slow proportion | None | None |
| **Automatic kinetic assay run?** | | Yes | No, end point assay | Yes |

Supplementary Table 9: Constructs used in this study

| Plasmid | Insert | Backbone/promoter | Insert sequence |
|---|---|---|---|
| rt256a | SARS-CoV-2 N-TEV-HaloTag-V5 | pHAGE/CMV | ATGTCTGATAATGGACCCCAAAATCAGCGAAATGCACCCCGCATTACGTTTGGTGGACCCTCAGATTCAACTGGC<br>AGTAACCAGAATGGAGAACGCAGTGGGGCGCGATCAAAACAACGTCGGCCCCAAGGTTTACCCAATAATACTGC<br>GTCTTGGTTCACCGCTCTCACTCAACATGGCAAGGAAGACCTTAAATTCCCTCGAGGACAAGGCGTTCCAATTAA<br>CACCAATAGCAGTCCAGATGACCAAATTGGCTACTACCGAAGAGCTACCAGACGAATTCGTGGTGGTGACGGTA<br>AAATGAAAGATCTCAGTCCAAGATGGTATTTCTACTACCTAGGAACTGGGCCAGAAGCTGGACTTCCCTATGGTG<br>CTAACAAAGACGGCATCATATGGGTTGCAACTGAGGGAGCCTTGAATACACCAAAAGATCACATTGGCACCCGC<br>AATCCTGCTAACAATGCTGCAATCGTGCTACAACTTCCTCAAGGAACAACATTGCCAAAAGGCTTCTACGCAGAA<br>GGGAGCAGAGGCGGCAGTCAAGCCTCTTCTCGTTCCTCATCACGTAGTCGCAACAGTTCAAGAAATTCAACTCC<br>AGGCAGCAGTAGGGGAACTTCTCCTGCTAGAATGGCTGGCAATGGCGGTGATGCTGCTCTTGCTTTGCTGCTGC<br>TTGACAGATTGAACCAGCTTGAGAGCAAAATGTCTGGTAAAGGCCAACAACAACAAGGCCAAACTGTCACTAAGA<br>AATCTGCTGCTGAGGCTTCTAAGAAGCCTCGGCAAAAACGTACTGCCACTAAAGCATACAATGTAACACAAGCTT<br>TCGGCAGACGTGGTCCAGAACAAACCCAAGGAAATTTTGGGGACCAGGAACTAATCAGACAAGGAACTGATTAC<br>AAACATTGGCCGCAAATTGCACAATTTGCCCCCAGCGCTTCAGCGTTCTTCGGAATGTCGCGCATTGGCATGGA<br>AGTCACACCTTCGGGAACGTGGTTGACCTACACAGGTGCCATCAAATTGGATGACAAAGATCCAAATTTCAAAGA<br>TCAAGTCATTTTGCTGAATAAGCATATTGACGCATACAAAACATTCCCACCAACAGAGCCTAAAAAGGACAAAAAG<br>AAGAAGGCTGATGAAACTCAAGCCTTACCGCAGAGACAGAAGAAACAGCAAACTGTGACTCTTCTTCCTGCTGCA<br>GATTTGGATGATTTCTCCAAACAATTGCAACAATCCATGAGCAGTGCTGACTCAACTCAGGCCGAGAACCTGTAC<br>TTCCAGTCTATGGCAGAAATCGGTACTGGCTTTCCATTCGACCCCCATTATGTGGAAGTCCTGGGCGAGCGCAT<br>GCACTACGTCGATGTTGGTCCGCGCGATGGCACCCCTGTGCTGTTCCTGCACGGTAACCCGACCTCCTCCTAC<br>GTGTGGCGCAACATCATCCCGCATGTTGCACCGACCCATCGCTGCATTGCTCCAGACCTGATCGGTATGGGCAA<br>ATCCGACAAACCAGACCTGGGTTATTTCTTCGACGACCACGTCCGCTTCATGGATGCCTTCATCGAAGCCCTGG<br>GTCTGGAAGAGGTCGTCCTGGTCATTCACGACTGGGGCTCCGCTCTGGGTTTCCACTGGGCCAAGCGCAATCC<br>AGAGCGCGTCAAAGGTATTGCATTTATGGAGTTCATCCGCCCTATCCCGACCTGGGACGAATGGCCAGAATTTG<br>CCCGCGAGACCTTCCAGGCCTTCCGCACCACCGACGTCGGCCGCAAGCTGATCATCGATCAGAACGTTTTTATC<br>GAGGGTACGCTGCCGATGGGTGTCGTCCGCCCGCTGACTGAAGTCGAGATGGACCATTACCGCGAGCCGTTCC<br>TGAATCCTGTTGACCGCGAGCCACTGTGGCGCTTCCCAAACGAGCTGCCAATCGCCGGTGAGCCAGCGAACAT<br>CGTCGCGCTGGTCGAAGAATACATGGACTGGCTGCACCAGTCCCCTGTCCCGAAGCTGCTGTTCTGGGGCACC<br>CCAGGCGTTCTGATCCCACCGGCCGAAGCCGCTCGCCTGGCCAAAAGCCTGCCTAACTGCAAGGCTGTGGACA |

| | | | |
|---|---|---|---|
| | | | TCGGCCCGGGTCTGAATCTGCTGCAAGAAGACAACCCGGACCTGATCGGCAGCGAGATCGCGCGCTGGCTGT<br>CGACGCTCGAGATTTCCGGCTCTAGTGGTGGAGGTAAGCCTATCCCTAACCCTCTCCTCGGTCTCGATTCTACG<br>TAA |
| rt189a | SARS-CoV-2 N-<br>SmBIT | pHAGE/HSVTK | ATGTCTGATAATGGACCCCAAAATCAGCGAAATGCACCCCGCATTACGTTTGGTGGACCCTCAGATTCAACTGGC<br>AGTAACCAGAATGGAGAACGCAGTGGGGCGCGATCAAAACAACGTCGGCCCCAAGGTTTACCCAATAATACTGC<br>GTCTTGGTTCACCGCTCTCACTCAACATGGCAAGGAAGACCTTAAATTCCCTCGAGGACAAGGCGTTCCAATTAA<br>CACCAATAGCAGTCCAGATGACCAAATTGGCTACTACCGAAGAGCTACCAGACGAATTCGTGGTGGTGACGGTA<br>AAATGAAAGATCTCAGTCCAAGATGGTATTTCTACTACCTAGGAACTGGGCCAGAAGCTGGACTTCCCTATGGTG<br>CTAACAAAGACGGCATCATATGGGTTGCAACTGAGGGAGCCTTGAATACACCAAAAGATCACATTGGCACCCGC<br>AATCCTGCTAACAATGCTGCAATCGTGCTACAACTTCCTCAAGGAACAACATTGCCAAAAGGCTTCTACGCAGAA<br>GGGAGCAGAGGCGGCAGTCAAGCCTCTTCTCGTTCCTCATCACGTAGTCGCAACAGTTCAAGAAATTCAACTCC<br>AGGCAGCAGTAGGGGAACTTCTCCTGCTAGAATGGCTGGCAATGGCGGTGATGCTGCTCTTGCTTTGCTGCTGC<br>TTGACAGATTGAACCAGCTTGAGAGCAAAATGTCTGGTAAAGGCCAACAACAACAAGGCCAAACTGTCACTAAGA<br>AATCTGCTGCTGAGGCTTCTAAGAAGCCTCGGCAAAAACGTACTGCCACTAAAGCATACAATGTAACACAAGCTT<br>TCGGCAGACGTGGTCCAGAACAAACCCAAGGAAATTTTGGGGACCAGGAACTAATCAGACAAGGAACTGATTAC<br>AAACATTGGCCGCAAATTGCACAATTTGCCCCCAGCGCTTCAGCGTTCTTCGGAATGTCGCGCATTGGCATGGA<br>AGTCACACCTTCGGGAACGTGGTTGACCTACACAGGTGCCATCAAATTGGATGACAAAGATCCAAATTTCAAAGA<br>TCAAGTCATTTTGCTGAATAAGCATATTGACGCATACAAAACATTCCCACCAACAGAGCCTAAAAAGGACAAAAAG<br>AAGAAGGCTGATGAAACTCAAGCCTTACCGCAGAGACAGAAGAAACAGCAAACTGTGACTCTTCTTCCTGCTGCA<br>GATTTGGATGATTTCTCCAAACAATTGCAACAATCCATGAGCAGTGCTGACTCAACTCAGGCCGGAGCTCAGGG<br>GAATTCTGGCTCGAGCGGTGGTGGCGGGAGCGGAGGTGGAGGGTCGTCAGGTGTGACCGGCTACCGGCTGTT<br>CGAGGAGATTCTGTAA |
| rt191a | SARS-CoV-2 N-<br>LgBIT | pHAGE/HSVTK | ATGTCTGATAATGGACCCCAAAATCAGCGAAATGCACCCCGCATTACGTTTGGTGGACCCTCAGATTCAACTGGC<br>AGTAACCAGAATGGAGAACGCAGTGGGGCGCGATCAAAACAACGTCGGCCCCAAGGTTTACCCAATAATACTGC<br>GTCTTGGTTCACCGCTCTCACTCAACATGGCAAGGAAGACCTTAAATTCCCTCGAGGACAAGGCGTTCCAATTAA<br>CACCAATAGCAGTCCAGATGACCAAATTGGCTACTACCGAAGAGCTACCAGACGAATTCGTGGTGGTGACGGTA<br>AAATGAAAGATCTCAGTCCAAGATGGTATTTCTACTACCTAGGAACTGGGCCAGAAGCTGGACTTCCCTATGGTG<br>CTAACAAAGACGGCATCATATGGGTTGCAACTGAGGGAGCCTTGAATACACCAAAAGATCACATTGGCACCCGC<br>AATCCTGCTAACAATGCTGCAATCGTGCTACAACTTCCTCAAGGAACAACATTGCCAAAAGGCTTCTACGCAGAA<br>GGGAGCAGAGGCGGCAGTCAAGCCTCTTCTCGTTCCTCATCACGTAGTCGCAACAGTTCAAGAAATTCAACTCC<br>AGGCAGCAGTAGGGGAACTTCTCCTGCTAGAATGGCTGGCAATGGCGGTGATGCTGCTCTTGCTTTGCTGCTGC |

| | | | |
|---|---|---|---|
| | | | TTGACAGATTGAACCAGCTTGAGAGCAAAATGTCTGGTAAAGGCCAACAACAACAAGGCCAAACTGTCACTAAGA<br>AATCTGCTGCTGAGGCTTCTAAGAAGCCTCGGCAAAAACGTACTGCCACTAAAGCATACAATGTAACACAAGCTT<br>TCGGCAGACGTGGTCCAGAACAAACCCAAGGAAATTTTGGGGACCAGGAACTAATCAGACAAGGAACTGATTAC<br>AAACATTGGCCGCAAATTGCACAATTTGCCCCCAGCGCTTCAGCGTTCTTCGGAATGTCGCGCATTGGCATGGA<br>AGTCACACCTTCGGGAACGTGGTTGACCTACACAGGTGCCATCAAATTGGATGACAAAGATCCAAATTTCAAAGA<br>TCAAGTCATTTTGCTGAATAAGCATATTGACGCATACAAAACATTCCCACCAACAGAGCCTAAAAAGGACAAAAAG<br>AAGAAGGCTGATGAAACTCAAGCCTTACCGCAGAGACAGAAGAAACAGCAAACTGTGACTCTTCTTCCTGCTGCA<br>GATTTGGATGATTTCTCCAAACAATTGCAACAATCCATGAGCAGTGCTGACTCAACTCAGGCCGGAGCTCAGGG<br>GAATTCTGGCTCGAGCGGTGGTGGCGGGAGCGGAGGTGGAGGGTCGTCAGGTGTCTTCACACTCGAAGATTTC<br>GTTGGGGACTGGGAACAGACAGCCGCCTACAACCTGGACCAAGTCCTTGAACAGGGAGGTGTGTCCAGTTTGC<br>TGCAGAATCTCGCCGTGTCCGTAACTCCGATCCAAAGGATTGTCCGGAGCGGTGAAAATGCCCTGAAGATCGAC<br>ATCCATGTCATCATCCCGTATGAAGGTCTGAGCGCCGACCAAATGGCCCAGATCGAAGAGGTGTTTAAGGTGGT<br>GTACCCTGTGGATGATCATCACTTTAAGGTGATCCTGCCCTATGGCACACTGGTAATCGACGGGGTTACGCCGA<br>ACATGCTGAACTATTTCGGACGGCCGTATGAAGGCATCGCCGTGTTCGACGGCAAAAAGATCACTGTAACAGGG<br>ACCCTGTGGAACGGCAACAAAATTATCGACGAGCGCCTGATCACCCCCGACGGCTCCATGCTGTTCCGAGTAAC<br>CATCAACAGCTAA |
| rt190a | HaloTag-SmBIT | pHAGE/HSVTK | ATGGCAGAAATCGGTACTGGCTTTCCATTCGACCCCCATTATGTGGAAGTCCTGGGCGAGCGCATGCACTACGT<br>CGATGTTGGTCCGCGCGATGGCACCCCTGTGCTGTTCCTGCACGGTAACCCGACCTCCTCCTACGTGTGGCGC<br>AACATCATCCCGCATGTTGCACCGACCCATCGCTGCATTGCTCCAGACCTGATCGGTATGGGCAAATCCGACAA<br>ACCAGACCTGGGTTATTTCTTCGACGACCACGTCCGCTTCATGGATGCCTTCATCGAAGCCCTGGGTCTGGAAG<br>AGGTCGTCCTGGTCATTCACGACTGGGGCTCCGCTCTGGGTTTCCACTGGGCCAAGCGCAATCCAGAGCGCGT<br>CAAAGGTATTGCATTTATGGAGTTCATCCGCCCTATCCCGACCTGGGACGAATGGCCAGAATTTGCCCGCGAGA<br>CCTTCCAGGCCTTCCGCACCACCGACGTCGGCCGCAAGCTGATCATCGATCAGAACGTTTTTATCGAGGGTACG<br>CTGCCGATGGGTGTCGTCCGCCCGCTGACTGAAGTCGAGATGGACCATTACCGCGAGCCGTTCCTGAATCCTG<br>TTGACCGCGAGCCACTGTGGCGCTTCCCAAACGAGCTGCCAATCGCCGGTGAGCCAGCGAACATCGTCGCGCT<br>GGTCGAAGAATACATGGACTGGCTGCACCAGTCCCCTGTCCCGAAGCTGCTGTTCTGGGGCACCCCAGGCGTT<br>CTGATCCCACCGGCCGAAGCCGCTCGCCTGGCCAAAAGCCTGCCTAACTGCAAGGCTGTGGACATCGGCCCG<br>GGTCTGAATCTGCTGCAAGAAGACAACCCGGACCTGATCGGCAGCGAGATCGCGCGCTGGCTGTCGACGCTCG<br>AGATTTCCGGCGTTTCTCAAGGCAGTTCAGGTGGTGGCGGGAGCGGAGGTGGAGGCTCGAGCGGTGTGACCG<br>GCTACCGGCTGTTCGAGGAGATTCTGTAA |

| | | | |
|---|---|---|---|
| rt163a | SARS-CoV-2 N-HaloTag | pHAGE/CMV | ATGTCTGATAATGGACCCCAAAATCAGCGAAATGCACCCCGCATTACGTTTGGTGGACCCTCAGATTCAACTGGC<br>AGTAACCAGAATGGAGAACGCAGTGGGGCGCGATCAAAACAACGTCGGCCCCAAGGTTTACCCAATAATACTGC<br>GTCTTGGTTCACCGCTCTCACTCAACATGGCAAGGAAGACCTTAAATTCCCTCGAGGACAAGGCGTTCCAATTAA<br>CACCAATAGCAGTCCAGATGACCAAATTGGCTACTACCGAAGAGCTACCAGACGAATTCGTGGTGGTGACGGTA<br>AAATGAAAGATCTCAGTCCAAGATGGTATTTCTACTACCTAGGAACTGGGCCAGAAGCTGGACTTCCCTATGGTG<br>CTAACAAAGACGGCATCATATGGGTTGCAACTGAGGGAGCCTTGAATACACCAAAAGATCACATTGGCACCCGC<br>AATCCTGCTAACAATGCTGCAATCGTGCTACAACTTCCTCAAGGAACAACATTGCCAAAAGGCTTCTACGCAGAA<br>GGGAGCAGAGGCGGCAGTCAAGCCTCTTCTCGTTCCTCATCACGTAGTCGCAACAGTTCAAGAAATTCAACTCC<br>AGGCAGCAGTAGGGGAACTTCTCCTGCTAGAATGGCTGGCAATGGCGGTGATGCTGCTCTTGCTTTGCTGCTGC<br>TTGACAGATTGAACCAGCTTGAGAGCAAAATGTCTGGTAAAGGCCAACAACAACAAGGCCAAACTGTCACTAAGA<br>AATCTGCTGCTGAGGCTTCTAAGAAGCCTCGGCAAAAACGTACTGCCACTAAAGCATACAATGTAACACAAGCTT<br>TCGGCAGACGTGGTCCAGAACAAACCCAAGGAAATTTTGGGGACCAGGAACTAATCAGACAAGGAACTGATTAC<br>AAACATTGGCCGCAAATTGCACAATTTGCCCCCAGCGCTTCAGCGTTCTTCGGAATGTCGCGCATTGGCATGGA<br>AGTCACACCTTCGGGAACGTGGTTGACCTACACAGGTGCCATCAAATTGGATGACAAAGATCCAAATTTCAAAGA<br>TCAAGTCATTTTGCTGAATAAGCATATTGACGCATACAAAACATTCCCACCAACAGAGCCTAAAAAGGACAAAAAG<br>AAGAAGGCTGATGAAACTCAAGCCTTACCGCAGAGACAGAAGAAACAGCAAACTGTGACTCTTCTTCCTGCTGCA<br>GATTTGGATGATTTCTCCAAACAATTGCAACAATCCATGAGCAGTGCTGACTCAACTCAGGCCAGCGGTGGTGGC<br>GGGAGCGGAGGTGGAGGGTCGTCAGGTATGGCAGAAATCGGTACTGGCTTTCCATTCGACCCCCATTATGTGG<br>AAGTCCTGGGCGAGCGCATGCACTACGTCGATGTTGGTCCGCGCGATGGCACCCCTGTGCTGTTCCTGCACGG<br>TAACCCGACCTCCTCCTACGTGTGGCGCAACATCATCCCGCATGTTGCACCGACCCATCGCTGCATTGCTCCAG<br>ACCTGATCGGTATGGGCAAATCCGACAAACCAGACCTGGGTTATTTCTTCGACGACCACGTCCGCTTCATGGAT<br>GCCTTCATCGAAGCCCTGGGTCTGGAAGAGGTCGTCCTGGTCATTCACGACTGGGGCTCCGCTCTGGGTTTCCA<br>CTGGGCCAAGCGCAATCCAGAGCGCGTCAAAGGTATTGCATTTATGGAGTTCATCCGCCCTATCCCGACCTGGG<br>ACGAATGGCCAGAATTTGCCCGCGAGACCTTCCAGGCCTTCCGCACCACCGACGTCGGCCGCAAGCTGATCAT<br>CGATCAGAACGTTTTTATCGAGGGTACGCTGCCGATGGGTGTCGTCCGCCCGCTGACTGAAGTCGAGATGGACC<br>ATTACCGCGAGCCGTTCCTGAATCCTGTTGACCGCGAGCCACTGTGGCGCTTCCCAAACGAGCTGCCAATCGC<br>CGGTGAGCCAGCGAACATCGTCGCGCTGGTCGAAGAATACATGGACTGGCTGCACCAGTCCCCTGTCCCGAAG<br>CTGCTGTTCTGGGGCACCCCAGGCGTTCTGATCCCACCGGCCGAAGCCGCTCGCCTGGCCAAAAGCCTGCCTA<br>ACTGCAAGGCTGTGGACATCGGCCCGGGTCTGAATCTGCTGCAAGAAGACAACCCGGACCTGATCGGCAGCGA<br>GATCGCGCGCTGGCTGTCGACGCTCGAGATTTCCGGCTAA |

| | | | |
|---|---|---|---|
| rt192a | SARS-CoV-2 N-NanoLuc | pHAGE/CMV | ATGTCTGATAATGGACCCCAAAATCAGCGAAATGCACCCCGCATTACGTTTGGTGGACCCTCAGATTCAACTGGC<br>AGTAACCAGAATGGAGAACGCAGTGGGGCGCGATCAAAACAACGTCGGCCCCAAGGTTTACCCAATAATACTGC<br>GTCTTGGTTCACCGCTCTCACTCAACATGGCAAGGAAGACCTTAAATTCCCTCGAGGACAAGGCGTTCCAATTAA<br>CACCAATAGCAGTCCAGATGACCAAATTGGCTACTACCGAAGAGCTACCAGACGAATTCGTGGTGGTGACGGTA<br>AAATGAAAGATCTCAGTCCAAGATGGTATTTCTACTACCTAGGAACTGGGCCAGAAGCTGGACTTCCCTATGGTG<br>CTAACAAAGACGGCATCATATGGGTTGCAACTGAGGGAGCCTTGAATACACCAAAAGATCACATTGGCACCCGC<br>AATCCTGCTAACAATGCTGCAATCGTGCTACAACTTCCTCAAGGAACAACATTGCCAAAAGGCTTCTACGCAGAA<br>GGGAGCAGAGGCGGCAGTCAAGCCTCTTCTCGTTCCTCATCACGTAGTCGCAACAGTTCAAGAAATTCAACTCC<br>AGGCAGCAGTAGGGGAACTTCTCCTGCTAGAATGGCTGGCAATGGCGGTGATGCTGCTCTTGCTTTGCTGCTGC<br>TTGACAGATTGAACCAGCTTGAGAGCAAAATGTCTGGTAAAGGCCAACAACAACAAGGCCAAACTGTCACTAAGA<br>AATCTGCTGCTGAGGCTTCTAAGAAGCCTCGGCAAAAACGTACTGCCACTAAAGCATACAATGTAACACAAGCTT<br>TCGGCAGACGTGGTCCAGAACAAACCCAAGGAAATTTTGGGGACCAGGAACTAATCAGACAAGGAACTGATTAC<br>AAACATTGGCCGCAAATTGCACAATTTGCCCCCAGCGCTTCAGCGTTCTTCGGAATGTCGCGCATTGGCATGGA<br>AGTCACACCTTCGGGAACGTGGTTGACCTACACAGGTGCCATCAAATTGGATGACAAAGATCCAAATTTCAAAGA<br>TCAAGTCATTTTGCTGAATAAGCATATTGACGCATACAAAACATTCCCACCAACAGAGCCTAAAAAGGACAAAAAG<br>AAGAAGGCTGATGAAACTCAAGCCTTACCGCAGAGACAGAAGAAACAGCAAACTGTGACTCTTCTTCCTGCTGCA<br>GATTTGGATGATTTCTCCAAACAATTGCAACAATCCATGAGCAGTGCTGACTCAACTCAGGCCAGCGGTGGTGGC<br>GGGAGCGGAGGTGGAGGGTCGTCAGGTATGGTCTTCACACTCGAAGATTTCGTTGGGGACTGGCGACAGACAG<br>CCGGCTACAACCTGGACCAAGTCCTTGAACAGGGAGGTGTGTCCAGTTTGTTTCAGAATCTCGGGGTGTCCGTA<br>ACTCCGATCCAAAGGATTGTCCTGAGCGGTGAAAATGGGCTGAAGATCGACATCCATGTCATCATCCCGTATGAA<br>GGTCTGAGCGGCGACCAAATGGGCCAGATCGAAAAAATTTTTAAGGTGGTGTACCCTGTGGATGATCATCACTTT<br>AAGGTGATCCTGCACTATGGCACACTGGTAATCGACGGGGTTACGCCGAACATGATCGACTATTTCGGACGGCC<br>GTATGAAGGCATCGCCGTGTTCGACGGCAAAAAGATCACTGTAACAGGGACCCTGTGGAACGGCAACAAAATTA<br>TCGACGAGCGCCTGATCAACCCCGACGGCTCCCTGCTGTTCCGAGTAACCATCAACGGAGTGACCGGCTGGCG<br>GCTGTGCGAACGCATTCTGGCGTAA |
| rt205a | CAHS2-SmBIT | pHAGE/HSVTK | ATGGAGGCAATGAATATGAATATCCCAAGGGATGCCATGTTTGTGCCCCCGCCAGAGTCTGAGCAAAACGGGTA<br>TCATGAAAAGTCTGAGGTTCAACAAACATCCTATATGCAGAGTCAGGTCAAGGTACCTCATTATAACTTCCCCACA<br>CCCTATTTTACGACTAGTTTTAGCGCCCAAGAGTTGTTGGGCGAAGGCTTTCAAGCTTCCATTTCTCGCATAAGCG<br>CCGTAACCGAAGACATGCAGAGTATGGAGATTCCAGAATTTGTCGAGGAAGCCAGACGAGATTATGCAGCAAAA<br>ACCCGCGAAAATGAGATGCTCGGCCAACAGTACGAAAAGGAATTGGAACGCAAAAGTGAGGCTTATCGAAAGCA<br>CCAAGAGGTTGAAGCTGATAAGATTCGCAAGGAATTGGAGAAACAACACATGAGAGACATCGAGTTCAGGAAAG |

| | | | |
|---|---|---|---|
| | | | AAATCGCGGAACTCGCTATTGAAAATCAAAAGCGGATGATCGACCTTGAATGCCGATATGCGAAGAAAGATATGG<br>ATCGAGAACGCACCAAAGTTCGAATGATGCTGGAACAACAAAAGTTTCATTCAGACATTCAGGTTAATCTTGACTC<br>CAGCGCTGCTGGGACAGAAAGTGGAGGGCATGTTGTAAGTCAAAGTGAGAAATTTACCGAGCGGAACCGCGAG<br>ATGAAAAGATCCGGAGGCGGCGGATCCACTAGAGGAGCTCAGGGGAATTCTGGCTCGAGCGGTGGTGGCGGG<br>AGCGGAGGTGGAGGGTCGTCAGGTGTGACCGGCTACCGGCTGTTCGAGGAGATTCTGTAA |
| rt206a | CAHS2-LgBIT | pHAGE/HSVTK | ATGGAGGCAATGAATATGAATATCCCAAGGGATGCCATGTTTGTGCCCCCGCCAGAGTCTGAGCAAAACGGGTA<br>TCATGAAAAGTCTGAGGTTCAACAAACATCCTATATGCAGAGTCAGGTCAAGGTACCTCATTATAACTTCCCCACA<br>CCCTATTTTACGACTAGTTTTAGCGCCCAAGAGTTGTTGGGCGAAGGCTTTCAAGCTTCCATTTCTCGCATAAGCG<br>CCGTAACCGAAGACATGCAGAGTATGGAGATTCCAGAATTTGTCGAGGAAGCCAGACGAGATTATGCAGCAAAA<br>ACCCGCGAAAATGAGATGCTCGGCCAACAGTACGAAAAGGAATTGGAACGCAAAAGTGAGGCTTATCGAAAGCA<br>CCAAGAGGTTGAAGCTGATAAGATTCGCAAGGAATTGGAGAAACAACACATGAGAGACATCGAGTTCAGGAAAG<br>AAATCGCGGAACTCGCTATTGAAAATCAAAAGCGGATGATCGACCTTGAATGCCGATATGCGAAGAAAGATATGG<br>ATCGAGAACGCACCAAAGTTCGAATGATGCTGGAACAACAAAAGTTTCATTCAGACATTCAGGTTAATCTTGACTC<br>CAGCGCTGCTGGGACAGAAAGTGGAGGGCATGTTGTAAGTCAAAGTGAGAAATTTACCGAGCGGAACCGCGAG<br>ATGAAAAGATCCGGAGGCGGCGGATCCACTAGAGGAGCTCAGGGGAATTCTGGCTCGAGCGGTGGTGGCGGG<br>AGCGGAGGTGGAGGGTCGTCAGGTGTCTTCACACTCGAAGATTTCGTTGGGGACTGGGAACAGACAGCCGCCT<br>ACAACCTGGACCAAGTCCTTGAACAGGGAGGTGTGTCCAGTTTGCTGCAGAATCTCGCCGTGTCCGTAACTCCG<br>ATCCAAAGGATTGTCCGGAGCGGTGAAAATGCCCTGAAGATCGACATCCATGTCATCATCCCGTATGAAGGTCT<br>GAGCGCCGACCAAATGGCCCAGATCGAAGAGGTGTTTAAGGTGGTGTACCCTGTGGATGATCATCACTTTAAGG<br>TGATCCTGCCCTATGGCACACTGGTAATCGACGGGGTTACGCCGAACATGCTGAACTATTTCGGACGGCCGTAT<br>GAAGGCATCGCCGTGTTCGACGGCAAAAAGATCACTGTAACAGGGACCCTGTGGAACGGCAACAAAATTATCGA<br>CGAGCGCCTGATCACCCCCGACGGCTCCATGCTGTTCCGAGTAACCATCAACAGCTAA |
| rt254a | CAHS2-Halo | pHAGE/CMV | ATGGAGGCAATGAATATGAATATCCCAAGGGATGCCATGTTTGTGCCCCCGCCAGAGTCTGAGCAAAACGGGTA<br>TCATGAAAAGTCTGAGGTTCAACAAACATCCTATATGCAGAGTCAGGTCAAGGTACCTCATTATAACTTCCCCACA<br>CCCTATTTTACGACTAGTTTTAGCGCCCAAGAGTTGTTGGGCGAAGGCTTTCAAGCTTCCATTTCTCGCATAAGCG<br>CCGTAACCGAAGACATGCAGAGTATGGAGATTCCAGAATTTGTCGAGGAAGCCAGACGAGATTATGCAGCAAAA<br>ACCCGCGAAAATGAGATGCTCGGCCAACAGTACGAAAAGGAATTGGAACGCAAAAGTGAGGCTTATCGAAAGCA<br>CCAAGAGGTTGAAGCTGATAAGATTCGCAAGGAATTGGAGAAACAACACATGAGAGACATCGAGTTCAGGAAAG<br>AAATCGCGGAACTCGCTATTGAAAATCAAAAGCGGATGATCGACCTTGAATGCCGATATGCGAAGAAAGATATGG<br>ATCGAGAACGCACCAAAGTTCGAATGATGCTGGAACAACAAAAGTTTCATTCAGACATTCAGGTTAATCTTGACTC<br>CAGCGCTGCTGGGACAGAAAGTGGAGGGCATGTTGTAAGTCAAAGTGAGAAATTTACCGAGCGGAACCGCGAG |

| | | | |
|---|---|---|---|
| | | | ATGAAAAGAAGCGGTGGTGGCGGGAGCGGAGGTGGAGGGTCGTCAGGTATGGCAGAAATCGGTACTGGCTTTC<br>CATTCGACCCCCATTATGTGGAAGTCCTGGGCGAGCGCATGCACTACGTCGATGTTGGTCCGCGCGATGGCAC<br>CCCTGTGCTGTTCCTGCACGGTAACCCGACCTCCTCCTACGTGTGGCGCAACATCATCCCGCATGTTGCACCGA<br>CCCATCGCTGCATTGCTCCAGACCTGATCGGTATGGGCAAATCCGACAAACCAGACCTGGGTTATTTCTTCGAC<br>GACCACGTCCGCTTCATGGATGCCTTCATCGAAGCCCTGGGTCTGGAAGAGGTCGTCCTGGTCATTCACGACTG<br>GGGCTCCGCTCTGGGTTTCCACTGGGCCAAGCGCAATCCAGAGCGCGTCAAAGGTATTGCATTTATGGAGTTCA<br>TCCGCCCTATCCCGACCTGGGACGAATGGCCAGAATTTGCCCGCGAGACCTTCCAGGCCTTCCGCACCACCGA<br>CGTCGGCCGCAAGCTGATCATCGATCAGAACGTTTTTATCGAGGGTACGCTGCCGATGGGTGTCGTCCGCCCG<br>CTGACTGAAGTCGAGATGGACCATTACCGCGAGCCGTTCCTGAATCCTGTTGACCGCGAGCCACTGTGGCGCTT<br>CCCAAACGAGCTGCCAATCGCCGGTGAGCCAGCGAACATCGTCGCGCTGGTCGAAGAATACATGGACTGGCTG<br>CACCAGTCCCCTGTCCCGAAGCTGCTGTTCTGGGGCACCCCAGGCGTTCTGATCCCACCGGCCGAAGCCGCTC<br>GCCTGGCCAAAAGCCTGCCTAACTGCAAGGCTGTGGACATCGGCCCGGGTCTGAATCTGCTGCAAGAAGACAA<br>CCCGGACCTGATCGGCAGCGAGATCGCGCGCTGGCTGTCGACGCTCGAGATTTCCGGCTAA |
| rt255a | CAHS2-NanoLuc | pHAGE/CMV | ATGGAGGCAATGAATATGAATATCCCAAGGGATGCCATGTTTGTGCCCCCGCCAGAGTCTGAGCAAAACGGGTA<br>TCATGAAAAGTCTGAGGTTCAACAAACATCCTATATGCAGAGTCAGGTCAAGGTACCTCATTATAACTTCCCCACA<br>CCCTATTTTACGACTAGTTTTAGCGCCCAAGAGTTGTTGGGCGAAGGCTTTCAAGCTTCCATTTCTCGCATAAGCG<br>CCGTAACCGAAGACATGCAGAGTATGGAGATTCCAGAATTTGTCGAGGAAGCCAGACGAGATTATGCAGCAAAA<br>ACCCGCGAAAATGAGATGCTCGGCCAACAGTACGAAAAGGAATTGGAACGCAAAAGTGAGGCTTATCGAAAGCA<br>CCAAGAGGTTGAAGCTGATAAGATTCGCAAGGAATTGGAGAAACAACACATGAGAGACATCGAGTTCAGGAAAG<br>AAATCGCGGAACTCGCTATTGAAAATCAAAAGCGGATGATCGACCTTGAATGCCGATATGCGAAGAAAGATATGG<br>ATCGAGAACGCACCAAAGTTCGAATGATGCTGGAACAACAAAAGTTTCATTCAGACATTCAGGTTAATCTTGACTC<br>CAGCGCTGCTGGGACAGAAAGTGGAGGGCATGTTGTAAGTCAAAGTGAGAAATTTACCGAGCGGAACCGCGAG<br>ATGAAAAGAAGCGGTGGTGGCGGGAGCGGAGGTGGAGGGTCGTCAGGTATGGTCTTCACACTCGAAGATTTCG<br>TTGGGGACTGGCGACAGACAGCCGGCTACAACCTGGACCAAGTCCTTGAACAGGGAGGTGTGTCCAGTTTGTTT<br>CAGAATCTCGGGGTGTCCGTAACTCCGATCCAAAGGATTGTCCTGAGCGGTGAAAATGGGCTGAAGATCGACAT<br>CCATGTCATCATCCCGTATGAAGGTCTGAGCGGCGACCAAATGGGCCAGATCGAAAAAATTTTTAAGGTGGTGTA<br>CCCTGTGGATGATCATCACTTTAAGGTGATCCTGCACTATGGCACACTGGTAATCGACGGGGTTACGCCGAACAT<br>GATCGACTATTTCGGACGGCCGTATGAAGGCATCGCCGTGTTCGACGGCAAAAAGATCACTGTAACAGGGACCC<br>TGTGGAACGGCAACAAAATTATCGACGAGCGCCTGATCAACCCCGACGGCTCCCTGCTGTTCCGAGTAACCATC<br>AACGGAGTGACCGGCTGGCGGCTGTGCGAACGCATTCTGGCGTAA |
